# De novo 3D models of SARS-CoV-2 RNA elements and small-molecule-binding RNAs to aid drug discovery

**DOI:** 10.1101/2020.04.14.041962

**Authors:** Ramya Rangan, Andrew M. Watkins, Jose Chacon, Wipapat Kladwang, Ivan N. Zheludev, Jill Townley, Mats Rynge, Gregory Thain, Rhiju Das

## Abstract

The rapid spread of COVID-19 is motivating development of antivirals targeting conserved SARS-CoV-2 molecular machinery. The SARS-CoV-2 genome includes conserved RNA elements that offer potential small-molecule drug targets, but most of their 3D structures have not been experimentally characterized. Here, we provide a compilation of chemical mapping data from our and other labs, secondary structure models, and 3D model ensembles based on Rosetta’s FARFAR2 algorithm for SARS-CoV-2 RNA regions including the individual stems SL1-8 in the extended 5’ UTR; the reverse complement of the 5’ UTR SL1-4; the frameshift stimulating element (FSE); and the extended pseudoknot, hypervariable region, and s2m of the 3’ UTR. For eleven of these elements (the stems in SL1-8, reverse complement of SL1-4, FSE, s2m, and 3’ UTR pseudoknot), modeling convergence supports the accuracy of predicted low energy states; subsequent cryo-EM characterization of the FSE confirms modeling accuracy. To aid efforts to discover small molecule RNA binders guided by computational models, we provide a second set of similarly prepared models for RNA riboswitches that bind small molecules. Both datasets (‘FARFAR2-SARS-CoV-2’, https://github.com/DasLab/FARFAR2-SARS-CoV-2; and ‘FARFAR2-Apo-Riboswitch’, at https://github.com/DasLab/FARFAR2-Apo-Riboswitch’) include up to 400 models for each RNA element, which may facilitate drug discovery approaches targeting dynamic ensembles of RNA molecules.

## Introduction

The COVID-19 outbreak has rapidly spread through the world, presenting an urgent need for therapeutics targeting the betacoronavirus SARS-CoV-2. RNA-targeting antivirals have potential to be effective against SARS-CoV-2, as the virus’s RNA genome harbors conserved regions predicted to have stable secondary structures (1,2) that have been verified by chemical probing (3–8), some of which have been shown to be essential for the life cycle of related betacoronaviruses (9). Efforts to identify small molecules that target stereotyped 3D RNA folds have advanced over recent years (10), making RNA structures like those in SARS-CoV-2 potentially attractive targets for small molecule drugs.

Several RNA regions in betacoronavirus genomes, including the 5’ UTR, the frameshift stimulating element (FSE), and 3’ UTR, feature RNA structures with likely functional importance. These regions include a series of five conserved stem-loops in the 5’ UTR, the frameshift stimulating element along with a proposed dimerized state, a pseudoknot in the 3’ UTR proposed to form two structures, and the hypervariable region in the 3’ UTR, which includes an absolutely conserved octanucleotide and the stem-loop II-like motif (‘s2m’). An NMR structure of stem-loop 2 in the 5’ UTR has been solved, adopting a canonical CUYG tetraloop fold (11). A crystal structure for s2m in the 3’ UTR has been solved for the original SARS virus, SARS-CoV-1 (12). Since reporting this work on the bioRxiv preprint server, structures for the frameshift stimulating element have been determined as an isolated RNA and in association with the ribosome through cryo-EM (13,14). Beyond these regions, however, 3D structures for RNA genome regions of SARS-CoV-2 or homologs have not been solved.

In advance of detailed experimental structural characterization, computational predictions for the 3D structural conformations adopted by conserved RNA elements may aid the search for RNA-targeting antivirals. Representative conformations from these RNA molecules’ structural ensembles can serve as starting points for virtual screening of small-molecules drug candidates. For example, a computational model for the FSE of SARS-CoV-1 was used in a virtual screen to discover the smallmolecule binder MTDB (15), and recently, SARS-CoV-2 models for 5’ UTR regions have been used for virtually docking small molecules (16). In other prior work by Stelzer, et al. (17), virtual screening of a library of compounds against an ensemble of modeled RNA structures led to the *de novo* discovery of a set of small molecules that bound a structured element in HIV-1 (the transactivation response element, TAR). Such work motivates our modeling of not just a single ‘native’ structure but an ensemble of states for SARS-CoV-2 RNA regions. As with HIV-1 TAR, many of the SARS-CoV-2 elements are unlikely to adopt a single conformation but instead sample conformations from a heterogeneous ensemble. Furthermore, transitions among these conformations may be implicated in the viral life cycle, as RNA genome regions change long-range contacts with other RNA elements or form interactions with viral and host proteins at different steps of replication, translation, and packaging. A possible therapeutic strategy is therefore to find drugs that stabilize an RNA element in a particular conformation incompatible with conformational changes and/or changing interactions with biological partners at different stages of the complete viral replication cycle. Consistent with this hypothesis, prior genetic selection and mutagenesis experiments stabilizing single folds for stemloops in the 5’ UTR and the pseudoknot in the 3′ UTR demonstrate that changes to these RNA elements’ structural ensembles can prove lethal for viral replication (18–20).

Here, we provide *de novo* modeled structure ensembles for conserved RNA elements in the SARS-CoV-2 genome obtained from Rosetta’s protocol for Fragment Assembly of RNA with Full-Atom Refinement, version 2 (FARFAR2) (21). These structures include *de novo* models for stem-loops 1 to 8 (SL1-8) in the extended 5’ UTR, the reverse complement of SL1-4 in the 5’ UTR, the FSE and its dimerized form, the 3′ UTR pseudoknot, and the 3′ UTR hypervariable region, along with homology models of SL2 and s2m. The use of Rosetta’s FARFAR2 is motivated by extensive testing: FARFAR2 has been benchmarked on all community-wide RNA-Puzzle modeling challenges to date (22–24), achieving accurate prediction of complex 3D RNA folds for ligand-binding riboswitches and aptamers, and producing models with 3–14 Å RMSD across six additional recent blind modeling challenges (21). For our SARS-CoV-2 study, the accuracy of our original *de novo* models for the FSE predicted in early 2020 has been validated by subsequent cryo-EM as well, as is described below. In addition to providing structural ensembles for SARS-CoV-2 RNA elements, we provide analogous FARFAR2 *de novo* and homology models for 10 riboswitch aptamers, providing a benchmark dataset for virtual screening approaches that make use of computational RNA models.

## Materials and Methods

### Chemical reactivity experiments

We collected chemical reactivity profiles for SL1-4 and SL2-6 of the 5’ UTR, the reverse complement of SL1-4, and the hypervariable region of the 3’ UTR. The DNA templates for the stem-loop 1-4 RNA were amplified from a gBlock sequence for the extended 5’ UTR, and the DNA template for the hyper-variable region was amplified from a gBlock sequence for the 3’ UTR. The SL2-6 construct was designed using the Primerize webserver (25) with built-in 5’ and 3’ “reference hairpins” for signal normalization flanking the region of interest and building using PCR assembly following the Primerize protocol (primers and gBlock sequences ordered from Integrated DNA Technologies, sequences in Table S5). For amplification off of gBlocks, primers were designed to add a Phi2.5 T7 RNA polymerase promoter sequence (26) (TTCTAATACGACTCACTATT) at the amplicon’s 5’ end and a 20 base-pair Tail2 sequence (AAAGAAACAACAACAACAAC) at its 3’ end. The PCR reactions contained 5 ng of gBlock DNA template, 2 μM of forward and reverse primer, 0.2 mM of dNTPs, 2 units of Phusion DNA polymerase, and 1X of HF buffer. The reactions were first denatured at 98 °C for 30 s. Then for 35 cycles, the samples were denatured at 98 °C for 10 sec, annealed at 64 °C for 30 sec, and extended at 72 C for 30 °C. This was followed by an incubation at 72 °C for 10 min for a final extension. Assembly products were verified for size via agarose gel electrophoresis and subsequently purified using Agencourt RNAClean XP beads. Purified DNA was quantified via NanoDrop (Thermo Scientific) and 8 pmol of purified DNA was then used for *in vitro* transcription with T7 TranscriptAid kits (Thermo Scientific). The resulting RNA was purified with Agencourt RNAClean XP beads supplemented with an additional 12% of PEG-8000 and quantified via NanoDrop. Owing to its longer length, the SL2-6 construct was subsequently size purified using a denaturing polyacrylamide gel (7M Urea, 1x TBE, hand-poured in Bio-Rad Criterion midi cassettes), loaded in 80% formamide, run at 18 Watts for 35 minutes following 1 hour of pre-running the gel prior to loading. A RiboRuler LR size standard (Thermo Scientific) was used. The correct-sized band was visualized using SyBr Gold (Invitrogen) and excised using a blue-light transilluminator, and the RNA was finally purified from the gel slice using a Zymo ZR-PAGE recovery kit.

For RNA modification, 1.2 pmol of RNA was denatured in 50 mM Na-HEPES pH 8.0 at 90 °C for 3 minutes and cooled at room temperature for 10 minutes. The RNA was then folded with the addition of MgCl_2_ to a final concentration of 10 mM in 15 μL and incubated at 50 °C for 30 minutes, then left at room temperature for 10 minutes. For chemical modification of folded RNA, fresh working stocks of 1-methyl-7-nitroisatoic anhydride (1M7) were prepared. For 1M7, 4.24 mg of 1M7 was dissolved in 1 mL of anhydrous DMSO. For a no-modification control reaction, 5 μL of RNase free H_2_O was added to 15 μL of folded RNA. Samples were incubated at room temperature for 15 minutes. Then, 5 μL of 5 M NaCl, 1.5 μL of oligo-dT Poly(A)Purist MAG beads (Ambion), and 0.065 pmol of 5’ Fluorescein (FAM)-labeled Tail2-A20 primer were added (sequence in Table S5), and the solution was mixed and incubated for 15 minutes. The magnetic beads were then pulled down by placing the mixture on a 96-post magnetic stand, washed twice with 100 μL of 70% EtOH, and air dried for 10 minutes before being resuspended in 2.5 μL RNase free H_2_O.

For cDNA synthesis, 2.5 μL resuspension of purified, polyA magnetic beads carrying chemically modified RNA was mixed with 2.5 μL of reverse transcription premix with SuperScript-III (Thermo Fisher). The reaction was incubated at 48 °C for 45 minutes. The RNA was then degraded by adding 5 μL of 0.4 M NaOH and incubating the mixture at 90 °C for 3 minutes. The degradation reaction was placed on ice and quickly quenched by the addition of 2 μL of an acid quench solution (1.4 M NaCl, 0.6 M HCl, and 1.3 M NaOAc). Bead-bound, FAM labeled cDNA was purified by magnetic bead separation, washed twice with 100 μL of 70% EtOH, and air-dried for 10 minutes. To elute the bound cDNA, the magnetic beads were resuspended in 10.0625 μL ROX/Hi-Di (0.0625 μL of ROX 350 ladder [Applied Biosystems] in 10 μL of Hi-Di formamide [Applied Biosystems]) and incubated at room temperature for 20 minutes. The resulting eluate was loaded onto capillary electrophoresis sequencers (ABI-3100 or ABI-3730) either on a local machine or through capillary electrophoresis (CE) services rendered by ELIM Biopharmaceuticals.

CE data was analyzed using the HiTRACE 2.0 package (https://github.com/ribokit/HiTRACE), following the recommended steps for sequence assignment, peak fitting, background subtraction of the no-modification control, correction for signal attenuation, and reactivity profile normalization.

### Eterna chemical mapping experiments

To probe mRNA structure, Eterna players designed 3030 sequences for the Eterna Roll Your Own Structure Lab, including regions of the SARS-CoV-2 5’ UTR, FSE, and 3’ UTR in Eterna Constructs 1-7 (Table S5). A DNA library for these constructs was synthesized by Genscript, with each construct 127 bases including the Phi2.5 T7 RNA polymerase promoter sequence (26) (TTCTAATACGACTCACTATT) and a 20 base-pair Tail2 sequence (AAAGAAACAACAACAACAAC) at its 3’ end.

This pool of DNA oligonucleotides (360 ng) was amplified by emulsion PCR with Phire Hot Start II DNA-Polymerase. An oil-surfactant mixture was prepared containing 80 μL of ABIL EM90, 1 μL of Triton X100 and 1919 μL of mineral oil. The oil phase was vortexed for 5 minutes and kept on ice for 30 minutes. An aqueous phase was prepared containing 1X Phire Hot Start II buffer, 0.2 mM dNTPs, 1.5 μL of Phire II DNA polymerase, 2 μL of the T7 promoter primer, 2 μM of the reverse complement of the Tail2 sequence, and 0.5 mg/ml of BSA in final volume of 75 μL. An emulsion was prepared in a 1.0 ml glass vial by first adding 300 μL of the oil-surfactant mixture into the glass vial, vortexing at 1000 rpm for 5 minutes, and then adding 10 μL of the aqueous phase every 10 seconds until the final emulsion volume was 350 μL. The emulsion was transferred into PCR tubes and PCR was performed by denaturing at 98 C for 30 seconds, cycling with 98 C for 10 seconds, 55 C for 10 seconds and 72 C for 30 seconds for 42 cycles, and extended at 72 C for 5 minutes. The PCR reaction was purified by adding 100 μL of mineral oil, vortexing, and centrifuging at 13,000 g for 10 minutes, and then discarding the oil phase. The PCR products were degreased with diethyl ether and ethyl acetate and incubated at 37 C for 5 minutes. The reaction volume was adjusted with H_2_O to 40 μL and then purified with 72 μL AMPure XP beads (Beckman Coulter), eluting into 20 μL of H_2_O.

DNA was transcribed with TranscriptAid T7 High Yield Transcription Kit (K0441), at 37 C for 3 hours, treated with DNAse-I for 30 minutes, and purified with AMPure XP beads with 40% PEG at a 7 to 3 ratio of beads to PEG. RNA was eluted with 25 μL of H_2_O. 15 pmol of RNA was added into 2 μL of 500 mM Na-HEPES, pH 8.0, denatured at 90 C for 3 minutes, and cooled down to room temperature for 10 minutes. 2 μL of 100 mM MgCl_2_ was added, and the reaction volume was brought to 15 μL with H_2_O. RNA was incubated at 50 C for 30 minutes. RNA was cooled down at room temperature for 20 minutes before being modified with 5 μL of 1M7 (8.48 mg/mL of DMSO) or left untreated for an untreated control sample. The reaction was left room temperature for 15 minutes, in final volume at 20 μL. The reaction was quenched with 5 μL of 500 mM Na-MES pH 6.0, the volume was adjusted to be 100 μL, and the reaction was purified with ethanol precipitation.

Reverse transcription was performed using SuperScript III RTase (Thermo Fisher). RNA was added into a reaction mix of 1X First strand buffer, 5 mM DTT, 0.8 mM dNTPs, and 0.6 μL of SS-III RTase (Thermo Fisher), and 1 μL of 0.25 μM primer (RTB000 and RTB001 in Table S5 for the no modification and 1M7 samples respectively, with these sequences adding an index sequence and Illumina adapter). The reaction volume was brought to 15 μL. The reaction was incubated at 48 C for 40 minutes and stopped by adding 5 μL of 0.4 M sodium hydroxide and heating the reaction at 90 C for 3 min, cooling the reaction on ice for 3 minutes, and neutralizing the reaction with 2 μL of an acid quench mix (2 mL of 5 M sodium chloride, 3 mL of 3 M sodium acetate, 2 mL of 2 M hydrochloric acid). cDNA was purified with Oligo C’ beads. Illumina adapters were ligated using Circ Ligase I (Lucigen) and linker pA-Adapt-Bp (Table S5), with ligation for 68 C for 2 hours, and the reaction was stopped at 80 C for 10 min. 10 μL of 5 M NaCl cDNA was added, and cDNA was purified with AMPure XP and eluted in 15 μL H2O. The ligated product was sequenced on a Miseq for 101 cycles for read 1 and 51 cycles for read 2. Sequencing data were analyzed using the MAPseeker software, freely available for non-commercial use at https://eternagame.org/about/software.

### Secondary structure modeling

Chemical reactivity from Manfredonia, et al. (8), Huston, et al. (6), and Sun, et al. (5) are publicly available at http://www.incarnatolab.com/datasets/SARS_Manfredonia_2020.php, http://www.github.com/pylelab/SARS-CoV-2_SHAPE_MaP_structure, and http://rasp.zhanglab.net respectively. DMS reactivity data from Lan, et al. (3) and and SHAPE reactivity data from Iserman, et al. (7) were obtained by request. We modeled RNA secondary structures using RNAstructure (27) guided by SHAPE or DMS reactivity data using default parameters, through MATLAB wrapper scripts available in the Biers package (https://github.com/ribokit/Biers).

### FARFAR2 3D modeling

We generated ensembles for SARS-CoV-2 RNA elements using Rosetta’s FARFAR2 protocol, providing a collection of models we term the **FARFAR2-SARS-CoV-2** dataset. Beginning with a sequence and secondary structure, FARFAR2 generates models through Monte Carlo substitutions of 3-residue fragments sampled from previously solved RNA structures, followed by refinement in a high-resolution physics-based free energy function, which models hydrogen bonding, solvation effects, nucleobase stacking, torsional preferences and other physical forces known to impact macromolecule structure (21). The models were created using the *rna_denovo* application in Rosetta 3.12 using default parameters for FARFAR2 (21). Rosetta is freely available for noncommercial use at https://www.rosettacommons.org.

For each system, we generated large model sets using the Stanford high performance computing cluster Sherlock and the Open Science Grid (28). For systems larger than 50 nucleotides, we clustered the 400 lowest energy structures with a 5 Å RMSD clustering radius, the procedure used for similarly sized systems in our recent FARFAR2 modeling benchmark (21). (Here and below, RMSD was computed as the all-heavy-atom RMSD between two models, as in all recent Rosetta work on RNA modeling.) For smaller systems, we clustered the 400 lowest energy structures with a 2 Å RMSD clustering radius, analogous to the procedure used for similarly sized systems in the FARFAR2 study. Clustering was achieved via the *rna_cluster* application used in the FARFAR2 and other Rosetta RNA studies, which iterates through unclustered structures from best to worst energy, either assigning them to an existing cluster (if the all-heavy-atom RMSD of the model is within the clustering radius of the cluster center) or starting a new cluster. We make available up to 50 models from each of the 10 lowest energy clusters in the resulting FARFAR2-SARS-CoV-2 data set to help efforts in virtual screening that take advantage of ensembles. We note that these models and their frequency in clusters are not necessarily an accurate representation of the thermodynamic ensemble obtained by the RNA due to biases in Rosetta FARFAR2 sampling and inaccuracies in the Rosetta all-atom free energy function. Nevertheless, the models offer a starting point of physically realistic conformations for virtual ligand screening and more sophisticated approaches to thermodynamic ensemble modeling.

For each RNA segment, we carried out Rosetta modeling using the secondary structure proposed in the literature (Table S1). We additionally considered experimentally derived secondary structures for each RNA region (Table S1). We carried out additional Rosetta modeling using each experimentally-derived secondary structure if the new secondary structure was substantially different from the original structure proposed in the literature (i.e., if it added or removed stems compared to the literature structure, or if it altered more than three base-pairs in any stem). Each 3D model collection reflects a single secondary structure for each RNA element; for constructs where more than one secondary structure has been proposed or predicted, we generated separate model collections, with the exception of the FSE for which we combined three closely related secondary structures.

Homology modeling with FARFAR2 for the 5’ UTR SL2 and the 3’ UTR stem-loop II-like motif (s2m) was carried out using the approach outlined in ref. (29). For the 5’ UTR SL2, PDB ID 2L6I (11) was used as a template for positions 45 to 59. For the 3’ UTR s2m, PDB ID 1XJR (12) was used as a template for positions 29728 to 29768; here, nucleotide numbering maps to the 3’ UTR secondary structure in Fig. 4.

### Quality assessment of models

Simulations that are sufficiently converged produce multiple occupancy clusters, which signal that FARFAR2 sampling is able to discover lowest energy states, as evaluated in Rosetta’s all-atom energy function. Runs with only single occupancy clusters would need more computer power to discover lowest energy states. In Table 1, we report the ‘E-gap’: the difference in Rosetta energy units (R.E.U.) for the best-scoring model in each cluster compared to the top-scoring model in the simulation overall. Rosetta energy functions have been fit such that R.E.U. estimate energies in kcal/mol (30), so E-gap values similar to or smaller than 4.0 indicate structures that are predicted to make up a significant fraction of the ground state ensemble and that may be trapped by small molecule drugs without a major cost in binding affinity.

**Table 1.** FARFAR2-SARS-CoV-2 models

| System/cluster | Length | Models generated | Model convergence (Å)<sup>a</sup> | Predicted minimum RMSD (Å)<sup>b</sup> | Percent of clusters with < 8.0 REU E-gap to lowest energy model<sup>c</sup> | Percent of clusters showing multiple occupancy<sup>d</sup> |
| --- | --- | --- | --- | --- | --- | --- |
| **5' UTR constructs** |  |  |  |  |  |  |
| 5' UTR (1–480) | 480 | 66011 | 50.9 | 44.92 ± 6.13 | 20% | 0% |
| 5' UTR stem-loop 1 (7–33) | 27 | 200000 | 1.83 | 5.17 ± 0.52 | 100% | 100% |
| 5' UTR stem-loop 2 (45–59) | 15 | 200000 | 2.39 | 5.63 ± 0.74 | 100% | 100% |
| 5' UTR stem-loop 3 (61–75) | 15 | 200000 | 2.58 | 5.78 ± 0.70 | 100% | 100% |
| 5' UTR stem-loop 4 (84–127) | 44 | 2018457 | 1.82 | 5.16 ± 0.49 | 100% | 80% |
| 5' UTR stem-loop 5 (148–295) | 148 | 2392320 | 18.99 | 19.07 ± 6.15 | 100% | 10% |
| 5' UTR stem-loop 5/6 (148–343) | 196 | 2020963 | 25.91 | 24.68 ± 6.27 | 50% | 10% |
| 5' UTR stem-loop 6 (302–343) | 42 | 200000 | 8.92 | 10.92 ± 2.21 | 100% | 100% |
| 5' UTR stem-loop 7 (349–394) | 27 | 200000 | 7.39 | 9.67 ± 1.90 | 20% | 100% |
| 5' UTR stem-loop 8 (407–478) | 72 | 4055322 | 6.86 | 9.24 ± 1.47 | 70% | 100% |
| **5' UTR reverse complement** |  |  |  |  |  |  |
| 5' UTR reverse complement stem-loops 1-4 (149–1) | 149 | 2031710 | 19.61 | 19.57 ± 4.37 | 90% | 0% |
| **Frameshift stimulating element constructs** |  |  |  |  |  |  |
| Frameshift stimulating element (13459–13546) | 88 | 390722 | 14.45 | 15.39 ± 3.09 | 80% | 80% |
| Suspected frameshift stimulating element dimer (13459–13546) | 176 | 23066 | 21.99 | 21.50 ± 4.08 | 30% | 0% |
| **3' UTR constructs** |  |  |  |  |  |  |
| 3' UTR beginning with bulged hairpin (29511–29871) | 361 | 11430 | 39.71 | 35.85 ± 5.51 | 20% | 0% |
| 3' UTR hypervariable region (29659–29852) | 194 | 28029 | 25.38 | 24.25 ± 4.18 | 80% | 0% |
| **3' UTR pseudoknot (29543–29665; 29846–29876)** | 158 | 1017205 | 21.93 | 21.45 ± 5.52 | 100% | 0% |
| **3' UTR pseudoknot fragment consisting of the pseudoknot (PK), P2, and P6 (29606–29665; 29846–29876)** | 95 | 1017205 | 10.24 | 11.99 ± 2.05 | 100% | 50% |
| **3' UTR BSL extended structure (29543–29665; 29846–29876)** | 158 | 1012716 | 24.04 | 23.16 ± 4.29 | 100% | 0% |
| **3' UTR stem-loop II-like motif, homology modeled from PDB ID: 1XJR(12) (29724–29773)** | 50 | 200000 | 6.95 | 9.32 ± 0.09 | 50% | 100% |
| **3' UTR stem-loop II-like motif, secondary structure based on NMR data from Wacker, et al.(35) (29724–29773)** | 50 | 500000 | 2.97 | 6.10 ± 0.75 | 100% | 10% |

<sup>a</sup>Mean pairwise all-heavy-atom RMSD between 10 lowest energy cluster centers discovered.
<sup>b</sup>Predicted RMSD to true structure.
<sup>c</sup>Rosetta all-atom free energy gap of cluster's lowest energy model compared to lowest energy model discovered in run. REU = Rosetta energy units, calibrated so that 1.0 corresponds approximately to 1 k<sub>B</sub>T.
<sup>d</sup>Percent of clusters with more than one cluster member. Clustering was carried out on top 400 models ranked by Rosetta all-atom free energy, based on 5.0 Å threshold, except for small RNAs (SL1-4, SL6-7, s2m), where 2.0 Å threshold was applied.

For each simulation, we additionally report a ‘convergence’ estimate in Table 1, estimated as the mean pairwise RMSD of the top 10 cluster centers predicted by FARFAR2. Prior work aiming at accurate prediction of single native crystal structures has demonstrated that convergence is a predictor for modeling accuracy (21,31,32), with prior tests suggesting that models that have 7.5 Å convergence or lower have mean single-structure prediction accuracy of at worst 10 Å, and models with 5 Å convergence or lower have single-structure prediction accuracy of at worst 8 Å (21). In this work, we are however not assuming that the RNA targets form a single ‘native’ structure. Instead, we take this convergence measure as a proxy for whether sampling may have been adequate to generate a useful model set. As a direct measure of the thoroughness of sampling, we also present the ‘occupancies’ of each of the top 10 clusters. Conformations sampled repeatedly in independent Rosetta-FARFAR2 runs (cluster membership greater than 1) indicate some level of convergence in sampling and those conformations are more likely to be realistic low-energy structures. In Figs. 1–4 below, we show cluster members as a cloud of translucent structures behind each cluster’s lowest energy conformation to visually convey the level of convergence; lack of such a cloud indicates a ‘singlet’ in which the cluster involves only one member. We include up to 50 representative top-scoring models in each cluster as the model collection for each RNA element, with structures available in the Github repository: https://github.com/DasLab/FARFAR2-SARS-CoV-2.

**Figure 1.**
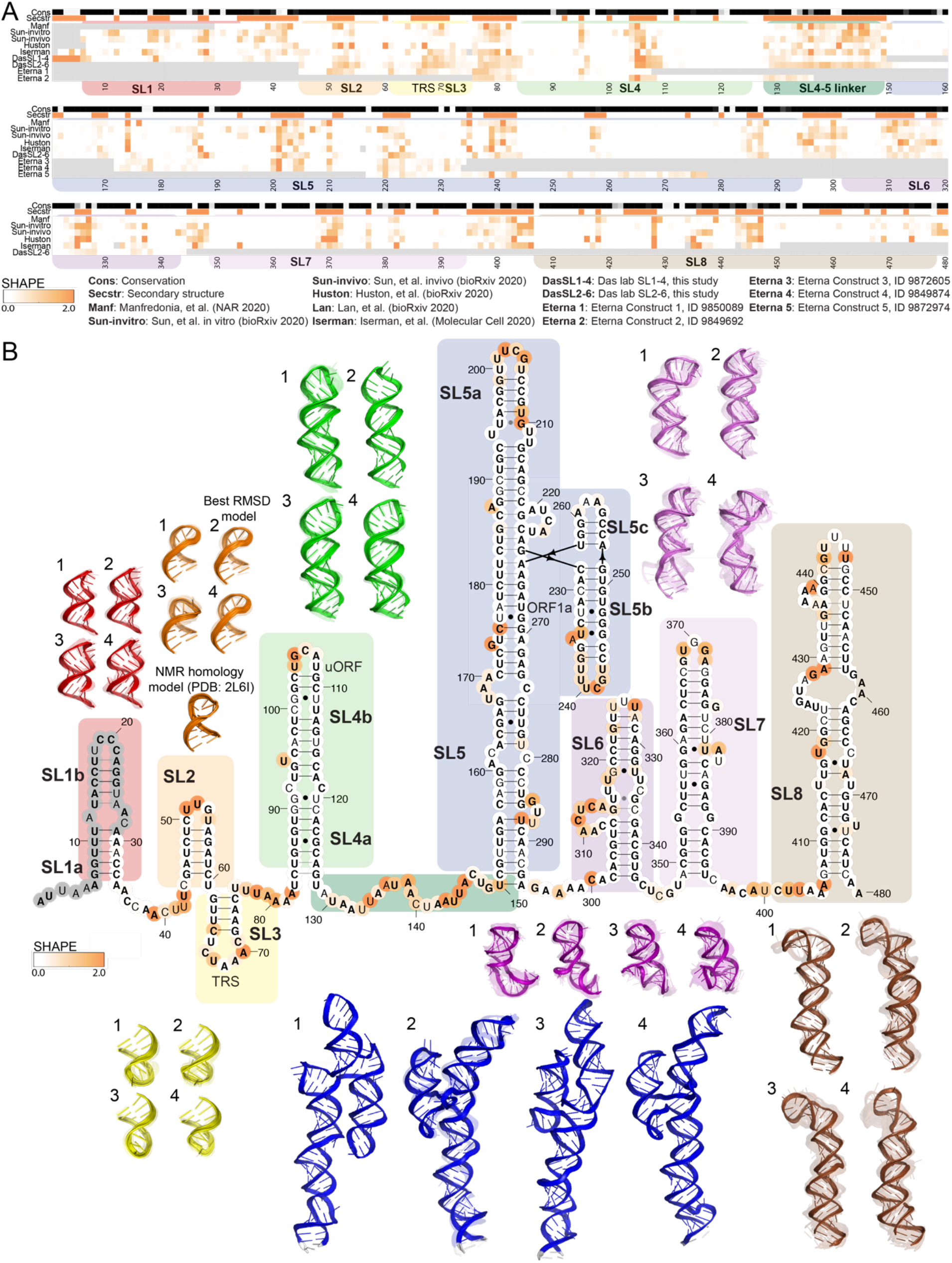
5’ UTR chemical reactivity, secondary structure, and 3D models. A) The heatmap compares chemical reactivity from recent publications probing SARS-CoV-2 RNA (3,5–8) along with reactivity data collected in this work (Das lab SL1-4 and Das lab SL2-6, and Eterna constructs 1-5). Gray values indicate no data, and reactivity increases from white to orange. The conservation track indicates the conservation percentage for each nucleotide across SARS-related species (54) from white (0% conserved) to black (100% conserved.) The secondary structure track is white in paired regions and orange in unpaired regions, following the secondary structure in panel B. Domains are indicated with coloring as follows: SL1 (red), SL2 (orange), SL3 (yellow), SL4 (light green), linker between SL4-SL5 (dark green), SL5 (blue), SL6 (purple), SL7 (light purple), and SL8 (brown). B) In bold are positions that are completely conserved across a set of SARS-related virus sequences (54). Base pairs that are not identified by the integrated DMS mapping and NMR analysis of Wacker, et al. (35) are shown in grey. Positions are colored according to their chemical reactivity in Manfredonia, et al. (8) Regions are boxed according to their coloring in 3D models. Top 4 clusters are depicted for SL1, SL2, SL3, SL4, SL5, SL6, SL7, and SL8. For SL2, a cluster derived from homology modeling to NMR structure 2L6I (11) is depicted, and the cluster with lowest RMSD from this NMR-derived structure is indicated. The top-scoring cluster member in each case is depicted with solid colors, and the top cluster members (up to 10) are depicted as transparent structures.

## Results

### Convergence of experimentally derived secondary structures across groups

RNA secondary structures are required to seed RNA 3D modeling. After our original modeling (33), several additional studies have been reported that constrain secondary structures through experimentally determined chemical reactivities of RNA segments *in vitro* or in cells (3,5–8,13). We made use of all currently published reactivity data along with data collected in our lab and on the Eterna project’s “cloud laboratory” pipeline to refine and update secondary structure models (Table S1; experimental methods described in Supplementary Methods). The SHAPE and DMS reactivity profiles for the 5’ UTR and beginning of the SARS-CoV-2 coding region, FSE, and 3’ UTR collected by different research groups across *in vitro* and *in vivo* probing conditions showed remarkable consistency across datasets (Fig. 1A, Fig. 3A, and Fig. 4A). The experimentally derived secondary structures were therefore similar across datasets (Table S1), with some exceptions including an extended region around the FSE and the hypervariable region of the 3’ UTR, noted below.

In addition to the growing wealth of chemical mapping data, recent NMR experiments integrated with DMS mapping determined the secondary structure for stem-loops in the 5’ UTR, the FSE, and regions of the 3’ UTR, largely confirming the secondary structures derived from chemical mapping experiments (34). In the 5’ UTR, these data support nearly all base pairs proposed previously in the literature, showing agreement with SHAPE and DMS reactivity experiments from recent studies. Additionally, the NMR data support the base pairs in the pseudoknot conformation for the FSE and the extended bulged stem-loop (BSL) conformation for the 3’ UTR pseudoknot, again agreeing with previously proposed structures. In these regions, the base pairs that are not seen in the NMR data are primarily terminal base pairs and are depicted in grey in the secondary structure diagrams in this manuscript. Notably, the SARS-CoV-2 s2m secondary structure determined by NMR differs from the secondary structure derived from homology modeling to the SARS-CoV-1 s2m crystal structure (1,12,34), providing a distinct secondary structure for Rosetta modeling (Table S1). The experimentally derived secondary structures and resulting 3D models are described in more detail for each probed segment below.

### Models of SARS-CoV-2 extended 5’ UTR

Fig. 1 presents models for the stem-loops that make up the extended 5’ UTR. Also called the 5’-proximal region, this region extends the 5’ UTR by ~200 residues to bracket potential structures that involve the beginning of the coding region. The secondary structure depicted in Fig. 1 is largely based on previous dissection of betacoronavirus secondary structures by several groups (9,35). More specifically, secondary structures for SL1-5 in the 5’ UTR are based on homology to prior betacoronaviruses, where these conserved stems have been confirmed through genetic experiments and sequence alignments in related betacoronaviruses. (We note for non-coronavirus researchers here that SL1, SL2, SL4, and SL5 have also been termed SLI, SLII, SLIII, and SLIV in an important set of studies (36–38).) Secondary structures for stems in the 5’ UTR were confirmed through predictions guided by chemical probing data from our group and five others (3,5–8), with all datasets predicting SL1-7 and generating structures for SL8 that had only minor variations across predictions (Table S1). These stems have been additionally validated by recent NMR experiments (34). Only terminal base-pairs in SL1, SL5, and SL6 are not identified by NMR, indicated in grey in Fig. 1.

Some of the stems in the extended 5’ UTR have structural preferences that have proven critical to the viral life cycle based on genetic experiments in other betacoronaviruses; these preferences have the potential to be altered through the binding of small-molecule drugs. Prior genetic selection experiments have demonstrated a preference for mutations that destabilize SL1 (red boxes in Fig. 1). The lower part of this stem must unpair to allow for the formation of a long-range RNA contact between the 5’ and 3’ UTRs (20). The stem must also presumably unfold to enable cap-dependent initiation of translation by the human ribosome. Recent work has found that SL1 is required for protecting SARS-CoV-2 mRNA from translation repression by SARS-CoV-2 protein nsp1, with SL1 likely binding to nsp1 as in SARS-CoV-1 (39,40). The loop in SL2 (orange boxes in Fig. 1) has sequence features across betacoronaviruses consistent with a U-turn conformation, and mutations that disrupt this structure are not viable, leading to a loss of sub-genomic RNA synthesis (18). SL3 (yellow boxes in Fig. 1) presents the transcription regulation sequence of the leader (TRS-L), which must be available to base-pair with TRS-B binding partners in the negative-strand viral genome to facilitate sub-genomic RNA synthesis (41). SL4 (light green boxes in Fig. 1) has a proposed role in directing the synthesis of subgenomic RNA in other betacoronaviruses (42), and also harbors an upstream open reading frame (uORF) across many betacoronaviruses. Though most RNAstructure predictions for the 5’ UTR maintained nucleotides between SL4 and SL5 as single-stranded (dark green boxes in Fig. 1), secondary structures based on the datasets of Sun, et al. (5) and Huston, et al. (6) point to the formation of an additional stem immediately 3’ of SL4. SL5 (blue boxes in Fig. 1) is a well-established domain that, in SARS-related viruses, has a long stem elaborated with a 4-way junction; this element has been proposed to harbor packaging signals, and it harbors the AUG start codon for the genome’s first gene product, the ORF1a/b polyprotein. Downstream of the AUG start codon, SL6 (purple boxes in Fig. 1) and SL7 (light purple boxes in Fig. 1) are predicted by all chemical mapping studies of the 5’ UTR (3,5–8), and are analogous to stems discovered to be important in bovine coronavirus but not yet functionally probed in SARS-related viruses (43). SL8 (brown boxes in Fig. 1) is also predicted by NMR and DMS reactivity experiments (3,34), though its function in SARS-CoV-2 is unknown.

We first produced models for the full extended 5’ UTR (over 1,000,000 FARFAR2 models generated on the Open Science Grid), with the top-scoring structures depicted in Fig. S2. With the current level of sampling, this and other lowest energy structures appear only once amongst our models (Table 1), reducing confidence that these models accurately capture lowest energy conformations. Nevertheless, the top-scoring models suggest the potential for compact RNA structures for the 5’ UTR mediated by potential tertiary contacts between stem-loops. While such tertiary contacts are of potential interest, additional experimental data would be needed to have confidence that any such collapsed states are well-defined low energy states and could act as therapeutic targets. It is also possible that the entire 5’ UTR may form a well-defined 3D arrangement when in complex with the ribosome, which would not be captured by our modeling. We therefore turned to smaller segments of the extended 5’ UTR for which Rosetta-FARFAR2 had a reasonable prospect of achieving convergent models.

For the shorter stem-loops in the 5’ UTR and initial stretch of the ORF1a/b coding region, including SL1, SL2, SL3, SL4, SL6, and SL7, we generated at least 200,000 FARFAR2 models. In Fig. 1, we depict the top four clusters for each stem-loop, and in Fig. S1 we depict the top ten clusters. We additionally modeled an extended SL4 construct that included the stem-loop immediately 3’ of SL4 predicted by Sun, et al. (5) and Huston, et al. (6) (Table S1, Fig. S1). As expected for these smaller RNA segments, all these stem-loops had excellent modeling convergence. Most clusters had occupancies of greater than 1, indicating that numerous independent *de novo* modeling trajectories resulted in conformations similar to within 2 Å RMSD. Furthermore, the mean pairwise RMSD of 10 lowest energy cluster centers (Table 1) approached 2.5 Å or better for SL1-4, suggesting that if a single dominant structure exists for these elements, our average model accuracy would be around 6 Å RMSD or better (Table 1). Nevertheless, SL1-3 have four or more clusters with E-gap values less than 1 R.E.U., suggesting the presence of many distinct structures at a 2 Å clustering radius. We propose that these clusters represent alternative structural targets that may be trapped by a small molecule without substantial energetic penalty.

In each of these RNA elements, stereotyped configurations of apical loops are modeled by Rosetta-FARFAR2 (Fig. 1). For SL2, a previous structure is available, determined by NMR for the SARS-CoV-1 sequence,^4^ providing a check on our modeling. We present a homology model of SL2 based on the SARS-CoV-1 sequence as an additional cluster in our data set (Fig. 1, see Supplementary Methods). Our *de novo* FARFAR2 models approach 3.1 Å RMSD to this homology-directed model, somewhat better than an accuracy estimate of RMSD of 5.6±0.7 Å from the native structure based on previously calibrated linear relationships and FARFAR2 modeling convergence of 2.4 Å (21).

For the larger stems of the extended 5’ UTR, we generated over 2,000,000 models per construct using the Open Science Grid (Fig 1, Fig S2). The SL5 element is a long stem-loop in all betacoronaviruses whose tip has been elaborated into a 4-way junction in SARS-CoV-2 and related subgroups (9). Due to the larger size and complexity of SL5, only one of the top 10 lowest energy models had another structure discovered within 5 Å RMSD among the top 400 lowest energy models (Table 1). Nevertheless, this structure did suggest the potential for drug binding pockets between helices that are brought into proximity by the four-way junction (Fig. 1). A joint simulation of SL5 and SL6 together did not produce convergence (Table 1). In the case of SL8, with over 4,000,000 models generated, we observed multiple occupancy clusters for all top ten clusters indicating sufficient convergence (Fig. 1, Table 1). Three clusters for SL8 have E-gap values below 4 R.E.U. (Table 1), suggesting that various alternate conformations for this stem-loop could be stabilized by interaction with small molecule drugs, especially in the terminal loop region which takes on unique conformations across top-scoring clusters.

### Reverse complement of SL1-4

We generated 3D models for the reverse complement of SL1-4, which may harbor secondary structures that bind to the viral replicase machinery during genome replication and transcription from the SARS-CoV-2 negative strand. For these models, we used a secondary structure derived from RNAstructure modeling guided by SHAPE data collected in this study (Fig. 2). The secondary structure includes four stem-loops: two short stems generated from the reverse complement of the 3’ strand of the 5’ UTR SL4, a longer branched stem-loop including a three-way junction and the full reverse-complements of the 5’ UTR SL2 and SL3, and a final short stem-loop comprising the reverse complement of the 5’ strand of the 5’ UTR SL1. With 750,000 models generated for this fragment, no clusters were generated with more than one member, indicating insufficient coverage. However, the three-way junction present in the central stem of this construct demonstrates tight packing and non-canonical interactions in some top-scoring models (Fig. 2B) and may show convergence with additional focused modeling.

**Figure 2.**
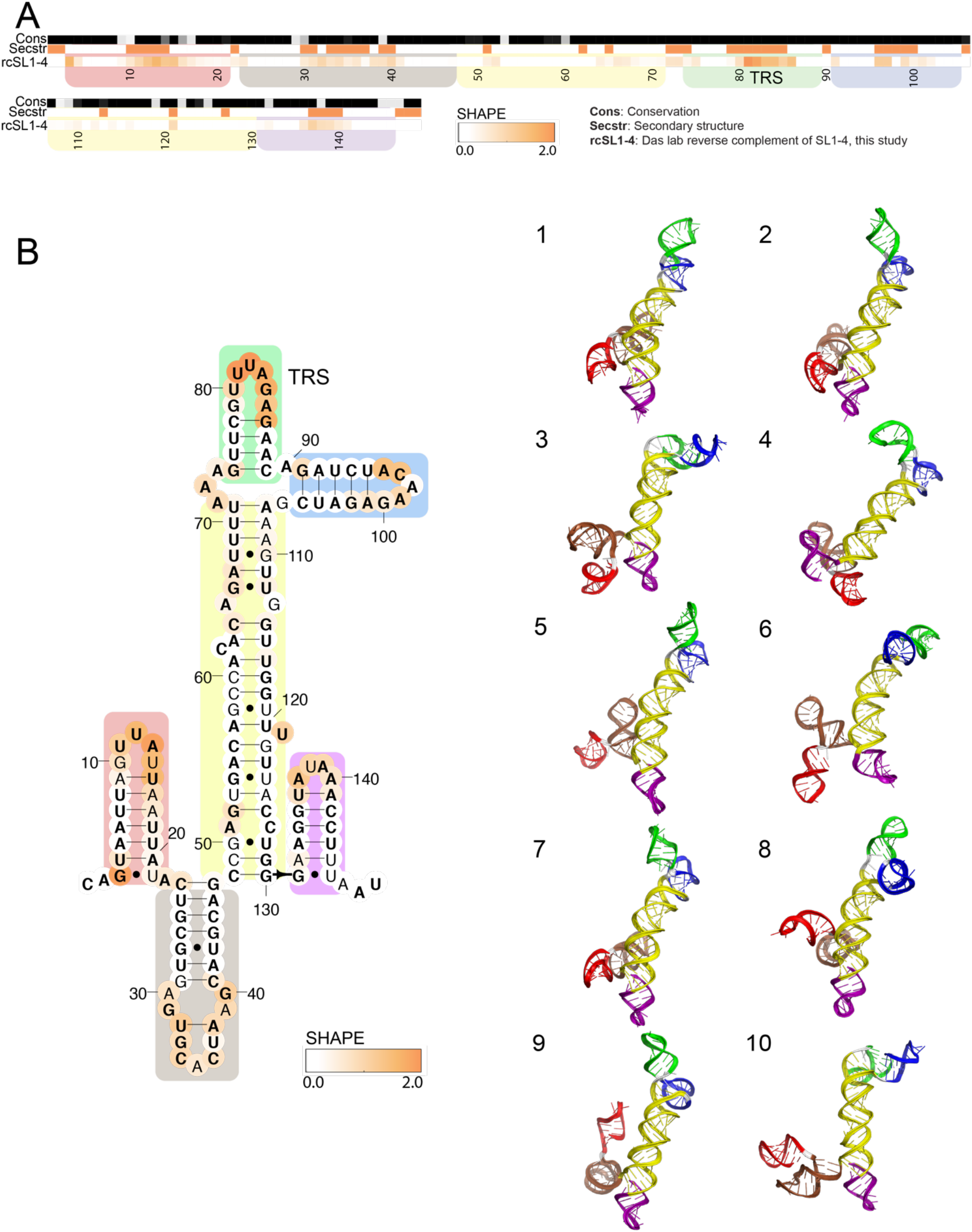
Chemical reactivity, secondary structure, and 3D models for the reverse complement of the 5’ UTR SL1-4. A) The heatmap depicts SHAPE reactivity from this work probing the reverse complement of the 5’ UTR SL1-4. Reactivity increases from white to orange. The conservation track indicates the conservation percentage for each nucleotide across SARS-related species (54) from white (0% conserved) to black (100% conserved.) The secondary structure track is white in paired regions and orange in unpaired regions, using the secondary structure as predicted by RNAstructure guided SHAPE data. Domains are indicated with colored boxes as follows. B) The secondary structure for the reverse complement of the 5’ UTR SL1-4 is depicted as used for FARFAR2 modeling. In bold are positions that are completely conserved across a set of SARS-related virus sequences.(54) Positions are colored according to their chemical reactivity shown in panel A). Regions are boxed according to their coloring in 3D models. 3D models for 10 clusters (all singleoccupancy) are depicted.

### Models of SARS-CoV-2 frameshift stimulating element

Fig. 3 presents Rosetta-FARFAR2 models for the SARS-CoV-2 frameshift stimulating element (FSE). The SARS-CoV-2 FSE pseudoknot structure has been shown to be critical for a (−1) ribosomal frameshifting event that leads to the production of ORF1a and ORF1b proteins from the same genomic region (Fig. 3) (44). The FSE has been the target of many recent structural characterization efforts. Recent chemical mapping studies have suggested alternative folds for the genomic region (Fig. 3 heatmap) (3,13), and cryo-EM structures for the FSE with and without the ribosome have been solved (13,14), providing independent checks for the *de novo* models produced in this work. Previously, a computer model enabled discovery of a small molecule ligand MTDB that is able to inhibit SARS-CoV-2 frameshifting, albeit with poor affinity (15,44–46), and a high-throughput screen has identified merafloxacin as a frameshifting inhibitor (47); these compounds reduce SARS-CoV-2 replication in Vero E6 cells, suggesting that targeting SARS-CoV-2 frameshifting rates may be a useful antiviral strategy (14,47).

**Figure 3.**
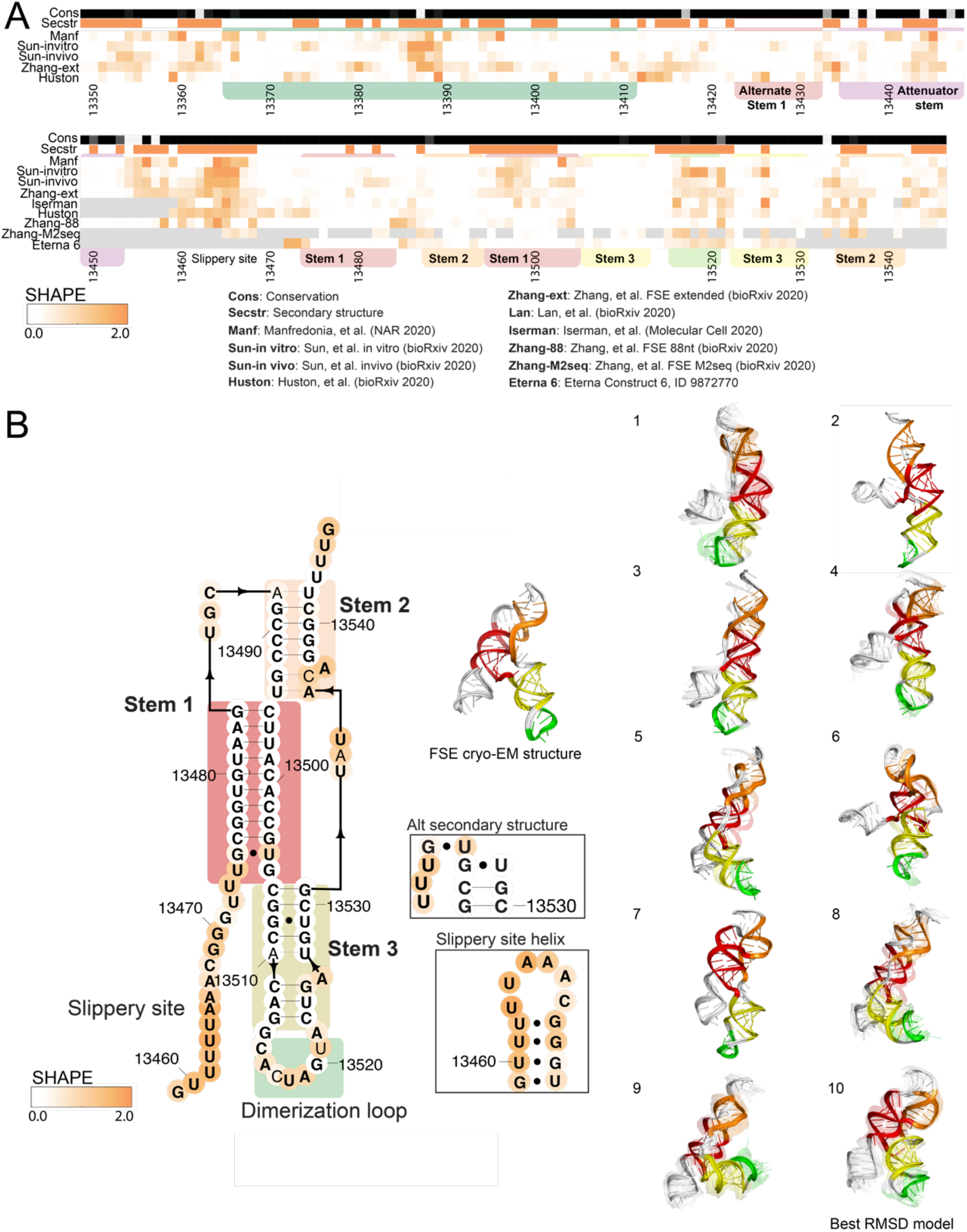
Frameshift stimulating element (FSE) chemical reactivity, secondary structure, and 3D models. A) The heatmap compares chemical reactivity from recent publications probing SARS-CoV-2 RNA (3,5–8,14), along with reactivity data collected in this work for a region within the FSE (Eterna construct 6). Gray values indicate no data, and reactivity increases from white to orange. The conservation track indicates the conservation percentage for each nucleotide across SARS-related species (54), from white (0% conserved) to black (100% conserved.) The secondary structure track is white in paired regions and orange in unpaired regions, using the secondary structure for the extended FSE is as predicted by RNAstructure guided SHAPE data (14). Domains are indicated with colored boxes as follows: Stem 1 (red), Stem 2 (orange), Stem 3 (yellow), and the dimerization loop (green). B) Frameshift stimulating element secondary structure, depicting alternate secondary structures used for FARFAR2 modeling. In bold are positions that are completely conserved across a set of SARS-related virus sequences (54). Base pairs that are not identified by the integrated DMS mapping and NMR analysis of Wacker, et al. (35) are shown in grey. Positions are colored according to their chemical reactivity when the 88-nt segment shown here was probed with SHAPE reagents (14). Regions are boxed according to their coloring in 3D models. 3D models for 10 frameshift stimulating element clusters are depicted. The top-scoring cluster member in each case is depicted with solid colors, and the top cluster members (up to 10) are depicted as transparent structures. The structure of the FSE as determined by cryo-EM in Zhang, et al. (14) is depicted, and the cluster center with lowest RMSD (9.9 Å) to this structure is indicated. Fig. S4 includes an alternate secondary structure and 3D models for the extended FSE including Alternate Stem 1.

To guide development of additional and more potent small molecules, we generated over 390,000 FARFAR2 models for the FSE. Of these, 100,000 models were generated separately with each of two similar secondary structures reported in the literature, which differ by a single base pair and include the single-stranded slippery site region (Fig. 3) (1,44). We noticed that many of the resulting top-scoring models contained a stem-loop surrounding the slippery-site sequence (Fig. 3). To focus sampling efforts on regions of the frame-shifting element beyond the slippery site, we generated 190,000 models with the slippery site helix pre-specified in the secondary structure, and with no base pairs formed with G13505. When all 390,000 models were considered together, the FSE simulation reached 14.4 Å convergence and yielded six clusters with more than one cluster member at a 5 Å RMSD clustering radius, with two clusters having E-gap values below 4 R.E.U. Models with all three of the assumed secondary structures produced low energies; they contribute to distinct but similar clusters. Despite the presence of various multiple occupancy clusters, compared to the smaller stem-loops modeled in the 5’ UTR, the FSE models were less converged. The majority of variation between top models arises from the 5’ slippery site sequence stem-loop, which adopts variable orientations in Rosetta-FARFAR2 simulations. The convergence of the pseudoknot region excluding this 5’ slippery sequence was tighter, 11.7 Å, suggesting that low-energy conformations of this pseudoknot element are captured by the current models.

After these *de novo* models for the FSE were generated, a cryo-EM structure of the frameshift stimulation element was solved by our group and collaborators (13). With 14.4 Å convergence between the top ten *de novo* models for the monomer FSE, we would expect an RMSD of 15.39±3.09 Å between the top FARFAR2 model and a single dominant native structure based on previously calibrated linear relationships (Table 1) (31,32). The best RMSD to the reported cryo-EM structure with lowest Rosetta energy was 9.88 Å. The RMSD between the best *de novo* model and the cryo-EM structure is better (lower) than the RMSD predicted by the convergence of the *de novo* models. Interestingly, the cryo-EM study showed clear density for the stem-loop surrounding the slippery-site sequence predicted from our *de novo* modeling. In addition, after these *de novo* models for the FSE were generated, the cryo-EM structure of the FSE on a mammalian ribosome was solved by Bhatt, et al. (14) We compared our *de novo* models to this structure for nucleotides 13471-13545, which form the pseudoknot just outside the mRNA channel entrance of the ribosome. For these positions, the top ten *de novo* models have a convergence of 12.23 Å, leading to an expected RMSD of 12.41±2.37 Å between the best of these ten models and the FSE structure on the ribosome. Indeed, the best RMSD between the *de novo* models and the FSE structure on the ribosome was 10.9 Å, falling within the expected range.

We additionally generated over 20,000 FARFAR2 models for a dimerized FSE, based on a proposal for dimerization through the loop of Stem 3 in SARS-CoV-1 (34,44,48). Modeling of the dimerized FSE did not converge on a well-defined 3D structure (each of the 10 lowest energy conformations were ‘singlets’ with no other conformations discovered within 5 Å RMSD; Table 1 and Fig. S3).

Recent chemical probing data on the FSE has suggested that in its genomic context, the FSE predominantly occupies an alternate conformation that does not include a pseudoknot (3,13).

Indeed, the 3’ strand of Stem 1 of the FSE appears reactive in most datasets (Fig. 3, right red box in heatmap), and RNAstructure predictions guided by chemical probing data from recent studies show a variety of structures when including ~100 nucleotides 5’ of the FSE (Table S1). While predictions using data from Manfredonia, et al. (8), Huston, et al. (6), and the *in vitro* probing condition from Sun, et al. (5) support formation of the FSE pseudoknot in this extended context, predictions using data collected in the other studies (3,5,7,13) summarized in the heatmap in Fig. 3 support an alternate pairing for the Stem 1 5’ strand with a region upstream of a previously characterized stem called the ‘attenuator hairpin’ (49). These differences may reflect the varied viral life cycle stages and probing conditions used to assay secondary structures in each of these studies. To begin exploring these alternate conformations, we generated FARFAR2 models using each of seven alternate secondary structures for the extended FSE (Table S1), generating 250,000 to 1,000,000 models for each case with the Open Science Grid. The top-scoring models did not reach convergence – each model reflected a structure not seen in other top scoring models – suggesting that the alternative secondary structures would not be associated with well-defined tertiary structures. Nevertheless, there is the potential for formation of an ensemble of heterogenous compact structures (Table S2, Fig. S4), and stems within this extended FSE are modeled with well-defined noncanonical features that may serve as targets for specific small-molecule binding. Indeed, recently, the attenuator hairpin has been targeted through a designed small molecule chimera that recruits RNase L to the stem (50).

### Models of SARS-CoV-2 3’ UTR

Fig. 4 presents models for structured regions of the 3’ UTR. The 3’ UTR includes a proposed switchlike pseudoknot element on the 5’ end, a hypervariable region, and the stem-loop II-like motif (Fig. 4A); these secondary structures are built based on homology to models of 3’ UTR’s of other betacoronaviruses (9,35). The 3’ UTR pseudoknot along with its mutually exclusive bulged stem-loop (BSL) structure have suggested functional roles in viral RNA synthesis, with mutations that destabilize either the pseudoknot or the stem-loop structure proving inviable in related betacoronaviruses (19). The structure of the stem-loop II-like motif resembles that of an rRNA loop, leading to its proposed role in recruiting host translation machinery (12).

**Figure 4.**
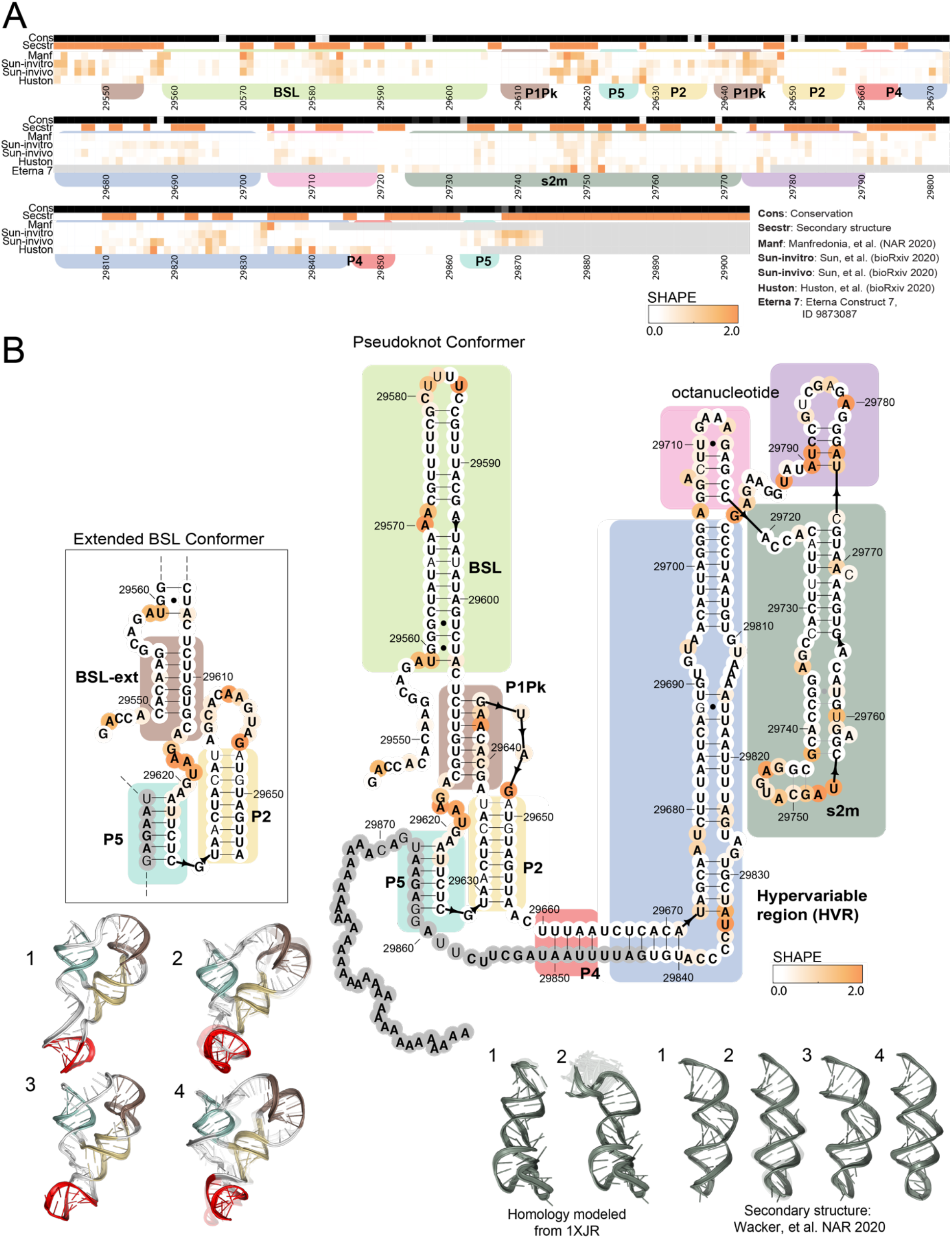
3’ UTR chemical reactivity, secondary structure, and 3D models. A) The heatmap compares chemical reactivity from recent publications probing SARS-CoV-2 RNA(3,5,6,8) along with reactivity data collected in this work. Gray values indicate no data, and reactivity increases from white to orange. The conservation track indicates the conservation percentage for each nucleotide across SARS-related species (54) from white (0% conserved) to black (100% conserved.) The secondary structure track is white in paired regions and orange in unpaired regions. Domains are indicated with colored boxes matching their coloring in 3D models. B) Positions are colored according to their chemical reactivity in Manfredonia, et al.(8) Regions are boxed according to their coloring in 3D models. In bold are positions that are completely conserved across a set of SARS-related virus sequences.(54) Base pairs that are not identified by the integrated DMS mapping and NMR analysis of Wacker, et al.(35) are shown in grey. 3D models are shown for the top 4 clusters for a segment containing the 3’ UTR pseudoknot, P2, and P5. 3D models are also shown for the top 4 clusters for the stem-loop II-like motif with models based on the NMR-derived secondary structure,(35) and the top 2 clusters for the stem-loop II-like motif, with models built based on homology to template structure 1XJR(12). The top-scoring cluster member in each case is depicted with solid colors, and the top cluster members (up to 10) are depicted as transparent structures.

We generated over 10,000 models for the full 3’ UTR. As with the models generated for the extended 5’ UTR, this simulation did not reach convergence (Table 1) but suggested the possibility of a heterogenous ensemble of compact RNA 3D structures (Fig. S5A). The 3’ UTR may form a well-defined 3D structure when complexed to other factors, such as the virus replicase/primase, but such a structure would not be captured with the RNA-only modeling carried out here.

We next turned to modeling two large subregions of the 3’ UTR: the 3’ UTR pseudoknot and the hyper-variable region. We built 1,000,000 models for the 3’ UTR pseudoknot on the Open Science Grid using the secondary structure depicted in Fig. 4, joining the 5’ and 3’ strands of the helix P4 with a tetraloop to generate a contiguous construct. Despite the large number of models generated for this simulation, we were not able to observe convergence due to the large size of this modeling case (Fig. S5B); none of the top 10 lowest energy conformations were similar to each other or to any of the other top 400 lowest energy conformations, as evaluated by RMSD with cutoff 5 Å (Table 1, Table S2). However, subregions of these modeling runs did achieve more convergence. For instance, the 3’ end of the models (positions 29606 to 29665 and 29842 to 29876) including the pseudoknot, P5, and P2 produced consistent compact conformations, with a convergence of 10.29 Å and various multiple occupancy clusters (Fig. 4, Table 1); this subregion may serve as a useful starting point for screening small-molecule RNA binders.

We additionally generated 2,000,000 models for the extended BSL structure that is mutually exclusive with the pseudoknot (Fig. 4 inset). When secondary structures for the 3’ UTR pseudoknot region were predicted using RNAstructure guided by recent chemical probing experiments, variants of the extended BSL structure were recovered in each case, suggesting that the pseudoknot secondary structure is not dominant in the conditions probed. The structure predicted by Huston, et al. (6) was the only predicted secondary structure that varied by more than three base-pairs from the BSL extended structure in Fig. 4 (Table S1). We therefore generated an additional 1,000,000 models for the 3’ UTR extended BSL with this secondary structure. Although these extended BSL simulations did not reach convergence (Figs. S5C and S6A), focused modeling of smaller intervals of the 3’ UTR may be fruitful for generating reasonable starting ensembles for virtual docking algorithms.

For the hyper-variable region (HVR), we generated over 25,000 models using the secondary structure depicted in Fig. 4, derived from Contrafold 2.0 (51) prediction and homology modeling to the SARS-CoV-1 s2m crystal structure (12). We observed that the RNAstructure predictions from chemical probing datasets yielded varying secondary structure predictions for this region, perhaps due to heterogeneity in this regions’ secondary structure ensemble (Table S1). We thus generated 1,000,000 additional models based on each experimentally derived secondary structure for this region (Table S1). However, modeling for the full HVR did not converge for any of these secondary structures (Fig. S6B-D). To further focus sampling, we turned to modeling the smaller SARS-CoV-1 stem-loop II-like motif (s2m) that is a part of the region. We used the crystal structure of the s2m to build over 200,000 homology models for the SARS-CoV-2 s2m with FARFAR2 (Fig. 4, see Supplementary Methods) (12,29). With near-identical sequences, the SARS-CoV-1 s2m template crystal structure is already a near-complete model for the SARS-CoV-2 domain, such that the FARFAR2 homology models for this region are highly converged, with the top ten models having an average RMSD of 0.21 Å. Recent chemical mapping experiments from this study and three other groups (5,6,8) along with NMR experiments (34) have predicted alternate secondary structures for the s2m (Table S1). We additionally generated s2m models for each of these secondary structures, producing converged model sets with multiple occupancy clusters at a 2Å radius (Fig. 4).

### Models of riboswitch aptamers as a benchmark for virtual drug screening methods

Use of the above SARS-CoV-2 RNA 3D models for virtual screening would be aided by a benchmark of analogous *de novo* models of RNA’s that are known to bind small molecules. Recent RNA-puzzles blind prediction trials have shown that FARFAR2 models in combination with conservation information enable manual identification of ligand binding sites in 3D models of bacterial ‘riboswitch’ aptamers (22–24). To guide use of FARFAR2 models for virtual screening, we have therefore collected a data set termed **FARFAR2-Apo-Riboswitch**, containing models of RNA elements that are known to bind small molecules, depicted in Fig. S7. These targets include binders for diverse ligands, from S-adenosyl methionine (SAM) to glycine, cobalamin, 5-hydroxytryptophan, the cyclic dinucleotides c-di-AMP (the ydaO riboswitch), the alarmone nucleotide ZMP (AICAR monophosphate), glutamine, and guanidinium. Each RNA was modeled without ligands present during fragment assembly or refinement, to mimic the protocols that would be used in virtual drug screening, in which modeling of *apo* RNA structures are used for computational docking of ligands.

Three of these model sets for SAM-I, SAM-I/IV, and SAM-IV made use of homology to previous riboswitch structures’ ligand binding sites (Homology, Fig. S7A-C), for historical reasons: the actual RNA-Puzzles challenges (or in the case of SAM-IV, an ‘unknown RFAM’ challenge for the RNA-Puzzles, involving tests based on cryoEM (52)) were posed at times in which crystal structures of riboswitch aptamers with homologous SAM binding sites were available. These model sets therefore serve as ‘positive controls’ for virtual drug screening protocols, which should be able to unambiguously identify homology-guided SAM binding sites as good aptamers for SAM.

The modeling also included riboswitch aptamer cases (*De novo*, Fig. S7D-J) in which the ligand binding sites were not modeled by homology, in closer analogy to virtual screening approaches that might make use of the FARFAR2-SARS-CoV-2 models. These models include cases such as the ydaO riboswitch, where modeling did not achieve a model closer than 10.0 Å RMSD to the RNA crystallized with two cyclic-diAMP ligands, perhaps owing to the large ligand and the substantial degree to which contacts with that ligand may organize the crystallized conformation. It will be interesting to see if these models still allow recognition of small molecule binding sites by computational methods. We provide in Table S4 additional metrics for these model sets, including the same RMSD convergence estimates, cluster occupancy, and E-gap numbers as for our FARFAR2-SARS-CoV-2 models as well as RMSD to experimentally determined ligand-bound structures. If these metrics correlate with the ability of virtual screening methods to discover known native ligands for these RNA elements, they will be useful in evaluating the likelihood of success of such methods in discovering molecules for the FARFAR2-SARS-CoV-2 models.

## Discussion

We have presented a collection of 3D models for elements comprising the extended 5’ UTR, reverse complement to the 5’ UTR, frameshift stimulating element, and 3’ UTR of the SARS-CoV-2 RNA genome. These models build on recent general advances in RNA 3D modeling, the ready availability of high performance computing, and a striking convergence of secondary structure modeling studies of the SARS-CoV-2 RNA from our and other laboratories. We hope that this FARFAR2-SARS-CoV-2 dataset provides a starting point for virtual screening approaches seeking compounds that stabilize individual SARS-CoV-2 RNA conformations, preventing access to functional conformations required for viral translation, replication, and packaging. Models for the 5’ UTR SL1-8 and the frameshift stimulating element appear to be especially promising candidates for small-molecule drug discovery. Modeling of these elements gave ensembles converging sufficiently to produce multiple representatives in clusters with 5 Å RMSD, and these elements all harbor sequences conserved across betacoronaviruses and structures having documented functional roles in the replication and/or translation of betacoronavirus genomes. The SL5 domain and the FSE present multi-helix 3D conformations with crevices and pockets that may be particularly amenable to small molecule targeting; validation of our FSE models through independent cryo-EM studies (13,14) further supports the accuracy of our models.

The structural ensembles presented here have a number of limitations. First, these ensembles are not true thermodynamic ensembles in that the structure occupancies do not necessarily reflect the underlying probabilities of occurrence for each conformational state. Additionally, some of the simulation ensembles described here did not achieve sufficient convergence to provide confidence in the resulting models (3’ UTR hypervariable region, extended 3’ UTR pseudoknot) – that is, independent modeling runs did not converge to similar low energy structures. It is possible that these RNA elements do not have well-defined 3D structures in solution unless bound tightly to partners such as the SARS-CoV-2 replicase complex. Alternatively, or in addition, our modeling methods and currently available computational power are not well-suited to regions of this size. While additional sampling may alleviate this problem, these regions have more *de novo* modeled positions than most prior FARFAR2 benchmark cases and may remain challenging for current *de novo* RNA modeling approaches.

Given these limitations in *de novo* modeling, we felt that it was important to provide analogous models of RNAs of known structure. Virtual screening approaches appear poised to make good use of computational models of RNA, but have so far made only limited use of *de novo* predicted models (15). To provide benchmark structural ensembles for such efforts, we have therefore used the same Rosetta-FARFAR2 modeling method and model selection procedure for a variety of RNA riboswitches with known small-molecule ligands. The resulting FARFAR2-Apo-riboswitch data set provides an opportunity for testing the accuracy of virtual screening approaches that use FARFAR2 model sets. With the current need for SARS-CoV-2 antiviral discovery, we believe this is an opportune time to explore and evaluate new approaches for virtual screening of small-molecule drug candidates that target structured RNA.

## Data availability

The supplementary file includes depictions of top-scoring cluster centers for the full extended 5’ UTR, the extended FSE with alternative secondary structures, the FSE dimer, the full 3’ UTR, the hypervariable region, and an extended 3’ UTR pseudoknot construct modeled with both the BSL and extended pseudoknot secondary structures. Chemical probing data collected in this study are available on RMDB (entries: FWSL14_UTR_0003 for SL1-4 in the 5’ UTR, FWSL26_UTR_0002 for SL2-6 in the 5’ UTR, RCSL14_UTR_0003 for the reverse complement of SL1-4 in the 5’ UTR, HVRS2M_UTR_0003 for the hyper-variable region in the 3’ UTR, and SHAPE_RYOS_0620 for the Eterna Roll Your Own Structure Lab). FARFAR2-SARS-CoV-2 models are included at https://github.com/DasLab/FARFAR2-SARS-CoV-2. FARFAR2-Apo-Riboswitch models are included at https://github.com/DasLab/FARFAR2-Apo-Riboswitch. Large model sets, comprising the top 5% of models for each simulation as ranked by Rosetta score, are included at the PURL repository https://purl.stanford.edu/pp620tj8748.

## Acknowledgments

We thank Silvi Rouskin, Cliff Zhang, Kevin Weeks, and Amy Gladfelter for providing chemical reactivity data upon request, and we thank Nenad Ban for providing coordinates for the frameshift stimulating element prior to deposition. We thank Sergey Lyskov and Jeff Gray for expedited access to the ROSIE cluster to pilot Rosetta modeling efforts. We thank Paul Berg, Jeff S. Glenn, Wah Chiu, and the Glenn and Chiu laboratories for illuminating discussions of the structures and functions of each presented element. The computing for this project was performed on the Sherlock cluster. We would like to thank Stanford University and the Stanford Research Computing Center for providing computational resources and support that contributed to these research results. This research was done using resources provided by the Open Science Grid (28).

