## Supplementary Information for "De novo 3D models of SARS-CoV-2 RNA elements and small-molecule-binding RNAs to aid drug discovery"

This supplemental information contains 7 Supplementary Figures and 4 Supplementary Tables.

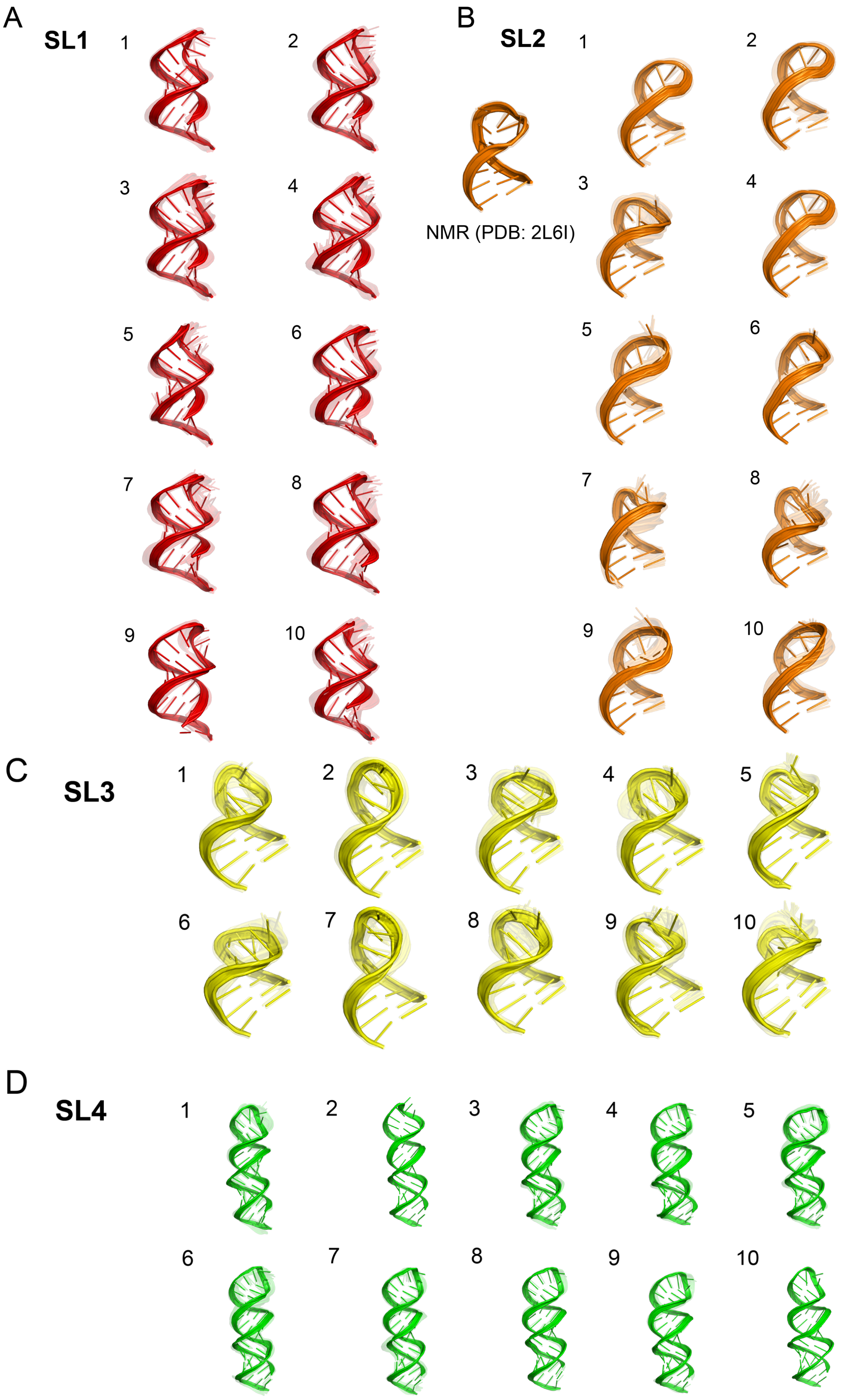
A
SL1
1
2
3
4
5
6
7
8
9
10
B
SL2
NMR (PDB: 2L6I)
1
2
3
4
5
6
7
8
9
10
C
SL3
1
2
3
4
5
6
7
8
9
10
D
SL4
1
2
3
4
5
6
7
8
9
10

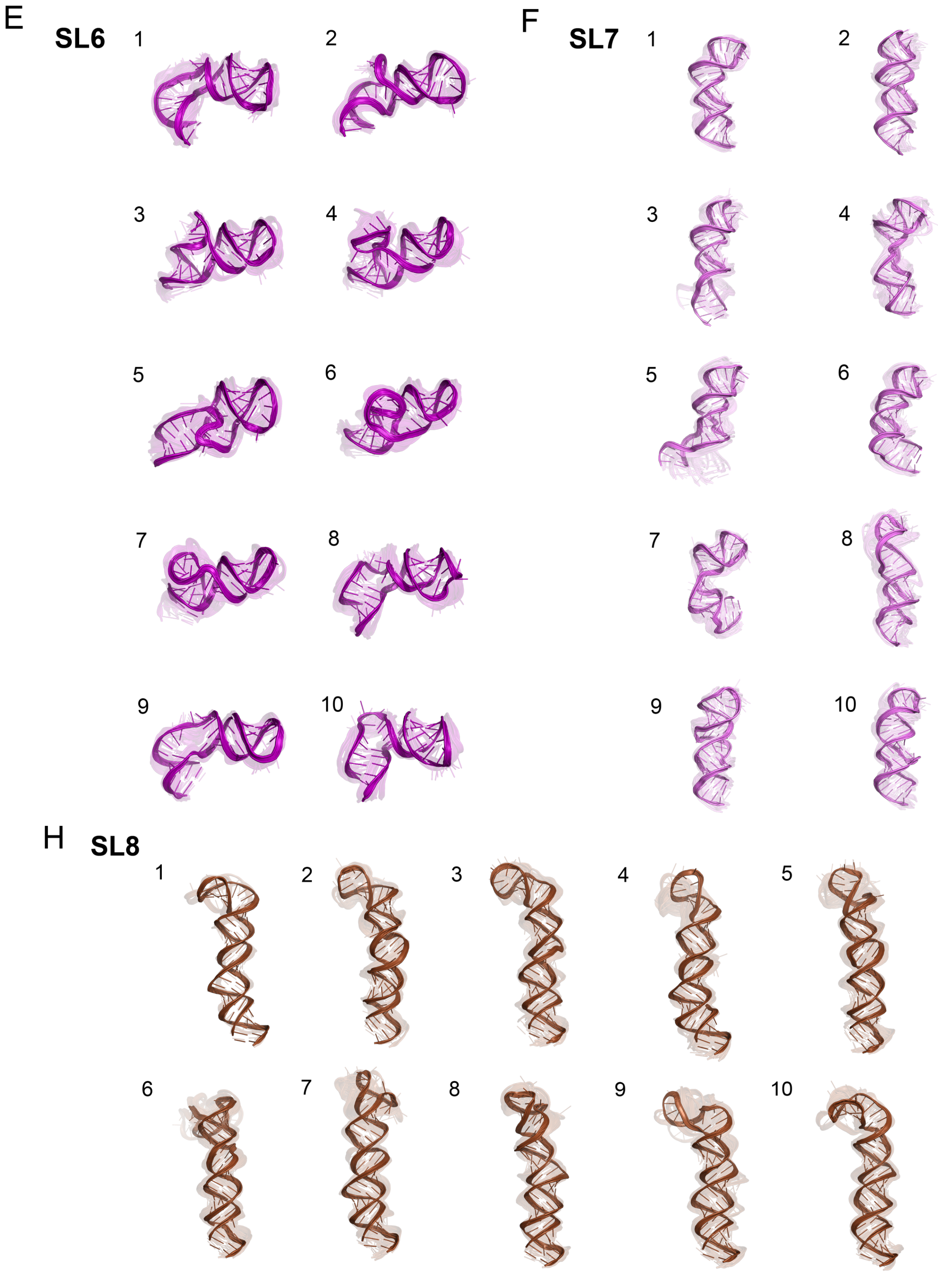
E
SL6
1
2
3
4
5
6
7
8
9
10
F
SL7
1
2
3
4
5
6
7
8
9
10
H
SL8
1
2
3
4
5
6
7
8
9
10

I

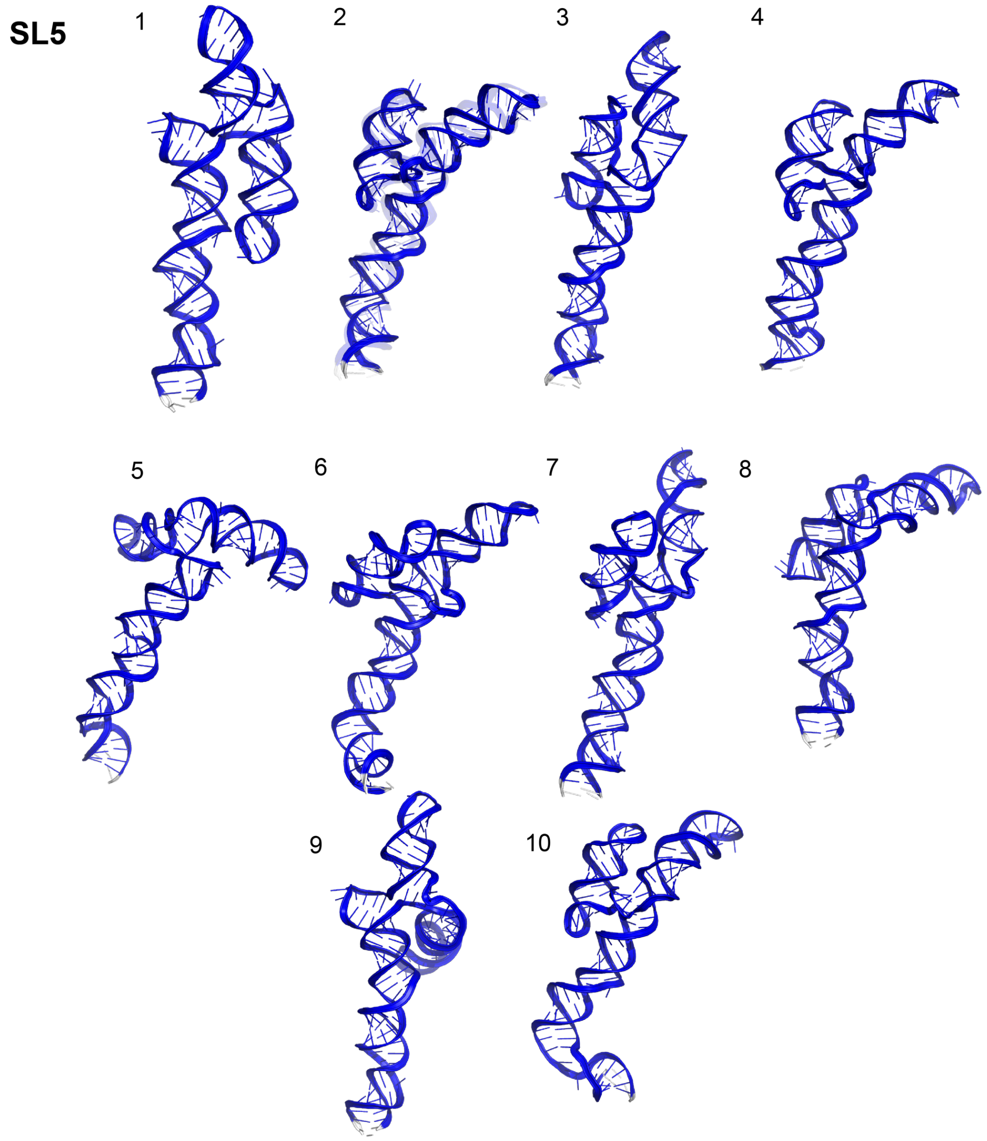


**Supplementary Figure 1.** *Models for the extended 5′ UTR.* Top 10 clusters are depicted for segments of the extended 5′ UTR: B) SL1, C) SL2, D) SL3, E) SL4, F) SL6, G) SL7, H) SL8, and I) SL5. Models are colored analogously to the secondary structure in Fig. 1.

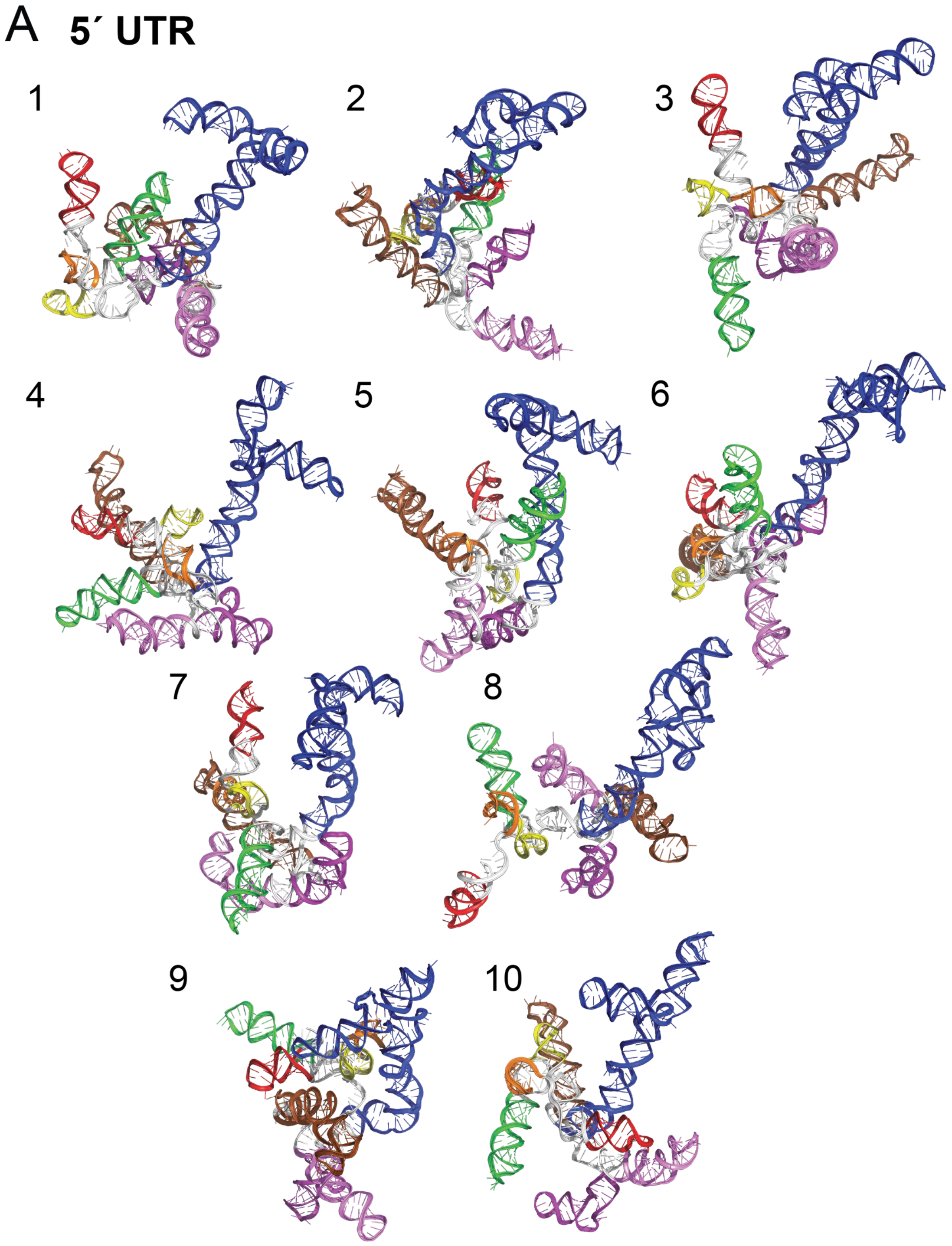
A
5´ UTR
1
2
3
4
5
6
7
8
9
10

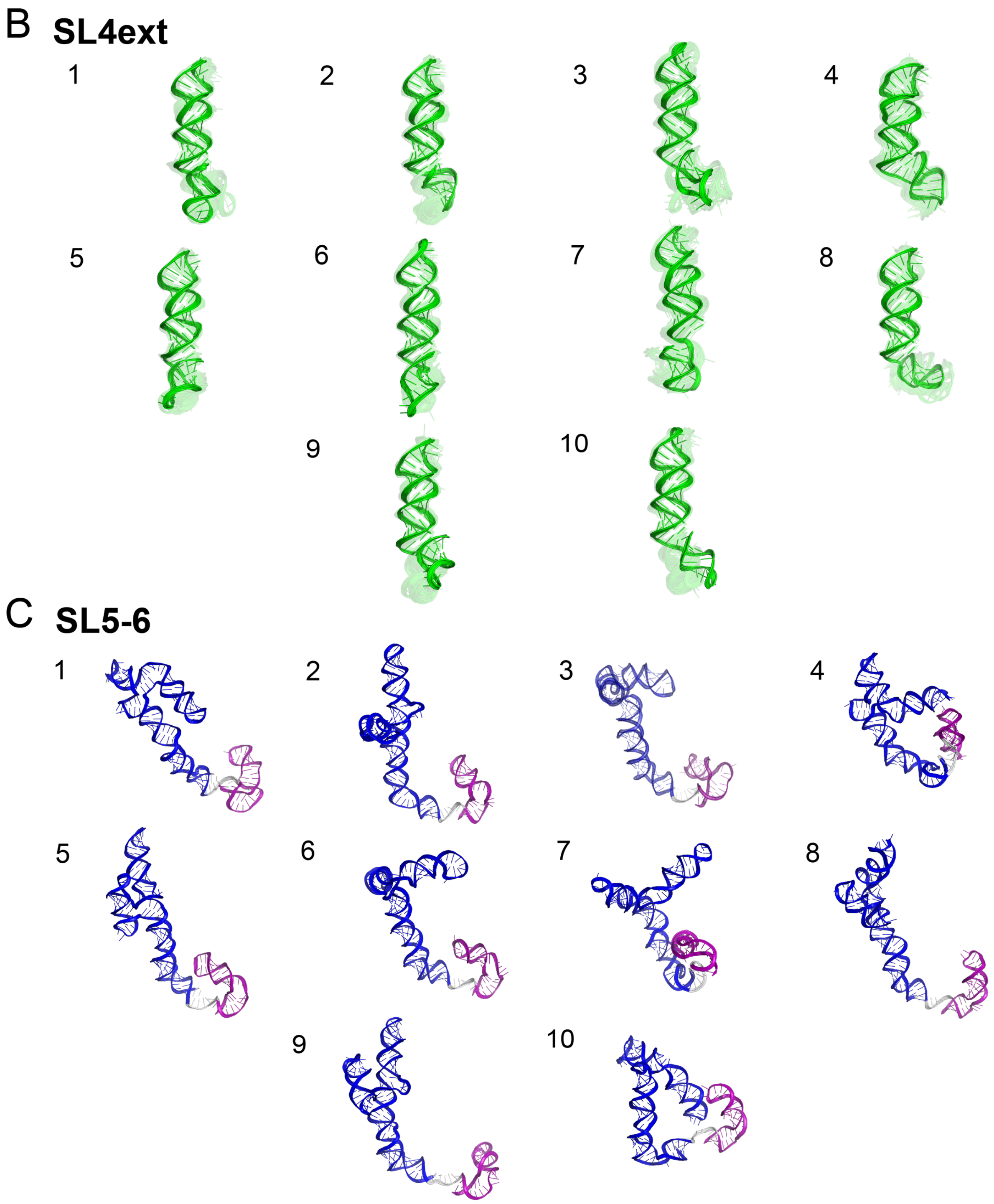


**Supplementary Figure 2.** *Models for the extended 5′ UTR, single occupancy clusters.* A) Top 10 clusters (all single-occupancy) are depicted for the extended 5′ UTR, colored analogously to the secondary structure in Fig. 1. B) Top 10 clusters for SL4 in the 5′ UTR along with a helix at the 3′ end predicted from some chemical mapping studies. C) Top 10 clusters for the SL5-6 in the 5′ UTR.

A

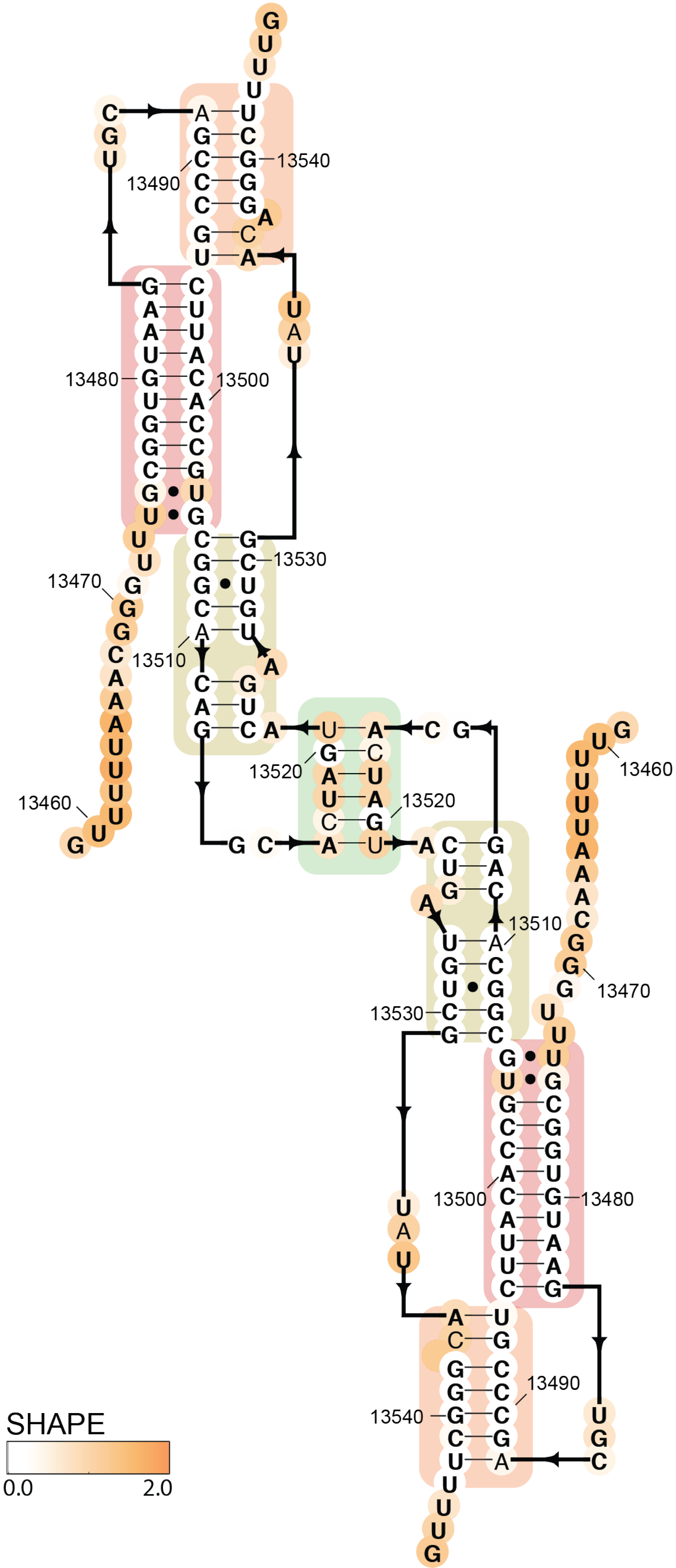
13490
13540
13480
13500
13470
13530
13510
13520
13520
13460
13460
13510
13470
13530
13500
13480
13490
13540
SHAPE
0.0
2.0

### B Frameshift stimulating element dimer

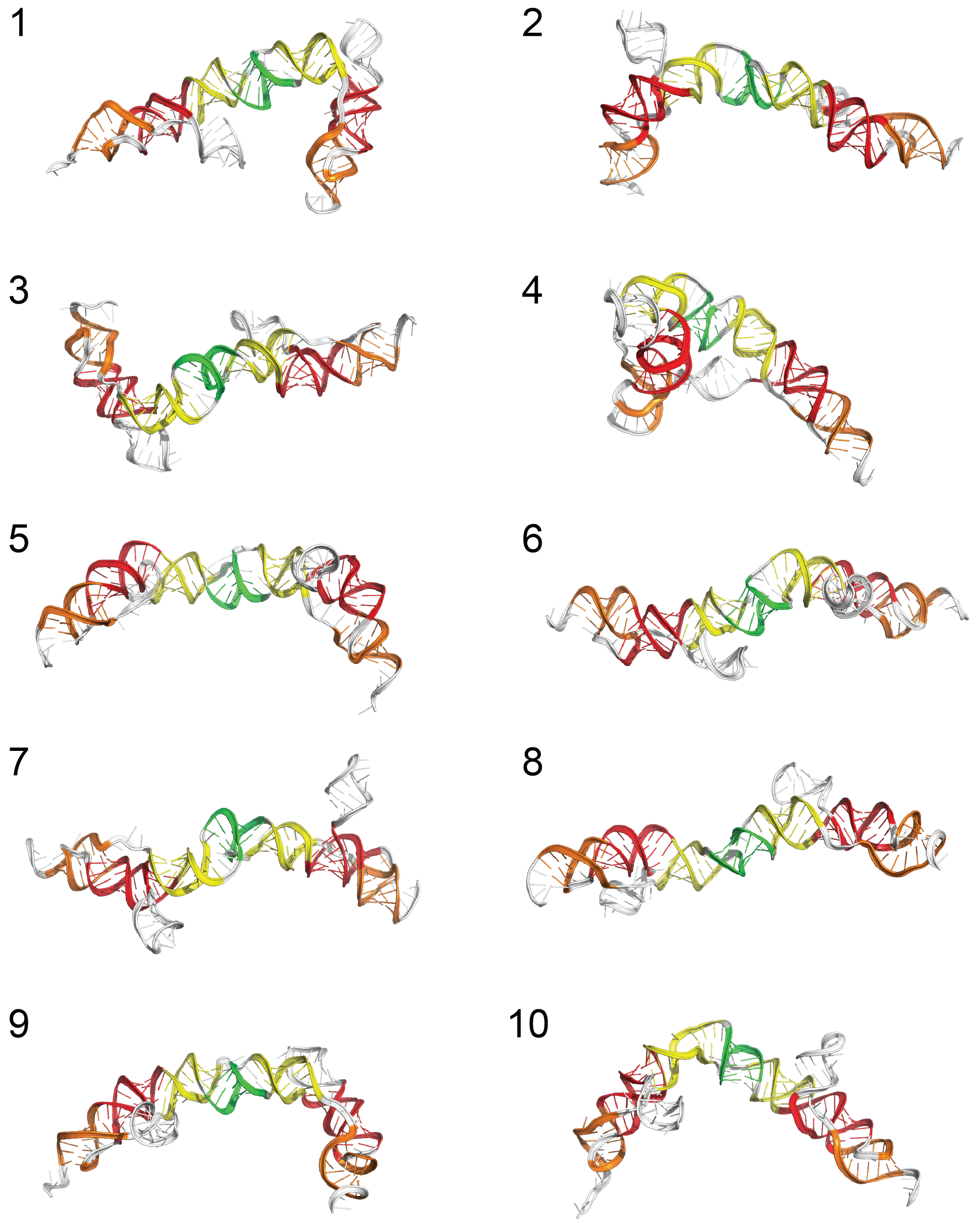


**Supplementary Figure 3.** *Models for the putative frameshift stimulation element dimer*. A) Secondary structure for the FSE dimer, generated from the secondary structure of the FSE in Fig. 3. B) Top 10 clusters (all single-occupancy) are depicted for the frameshift stimulating element dimer, colored analogously to the secondary structure in Fig. 3.

A

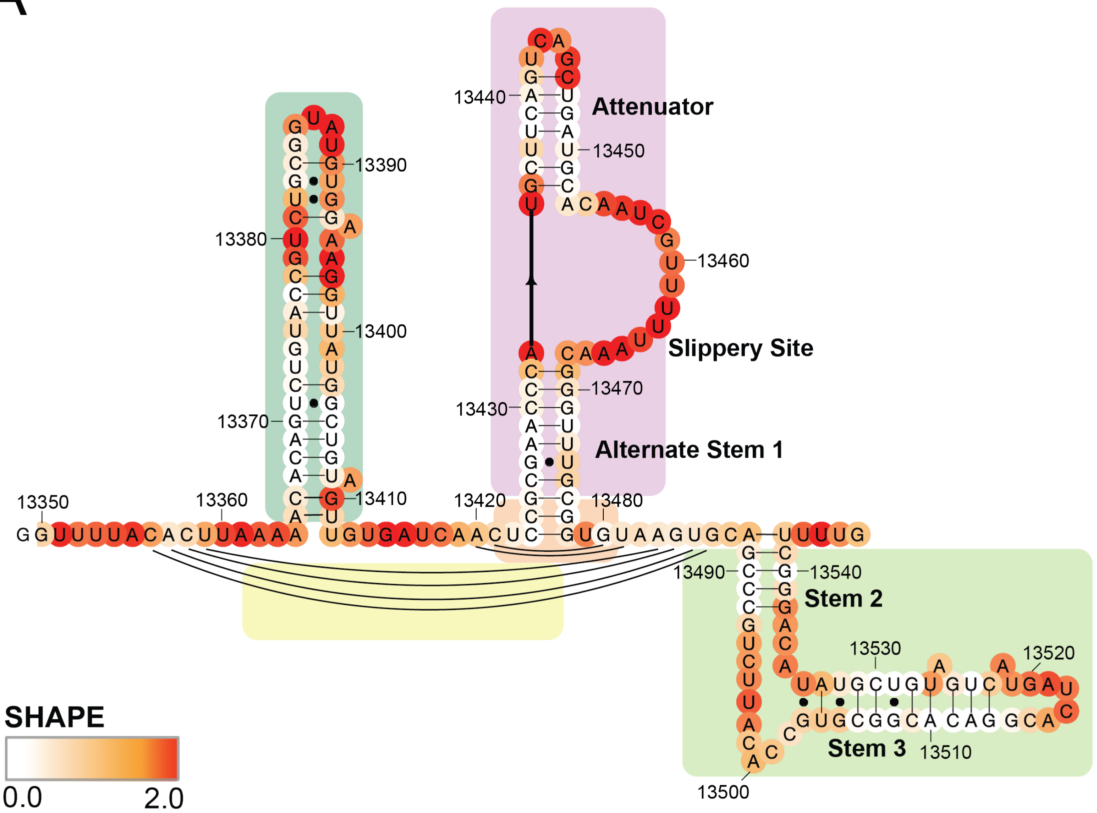

Attenuator
Slippery Site
Alternate Stem 1
Stem 2
Stem 3
13350
13360
13370
13380
13390
13400
13410
13420
13430
13440
13450
13460
13470
13480
13490
13500
13510
13520
13530
13540
SHAPE
0.0
2.0

### Extended frameshift stimulating element

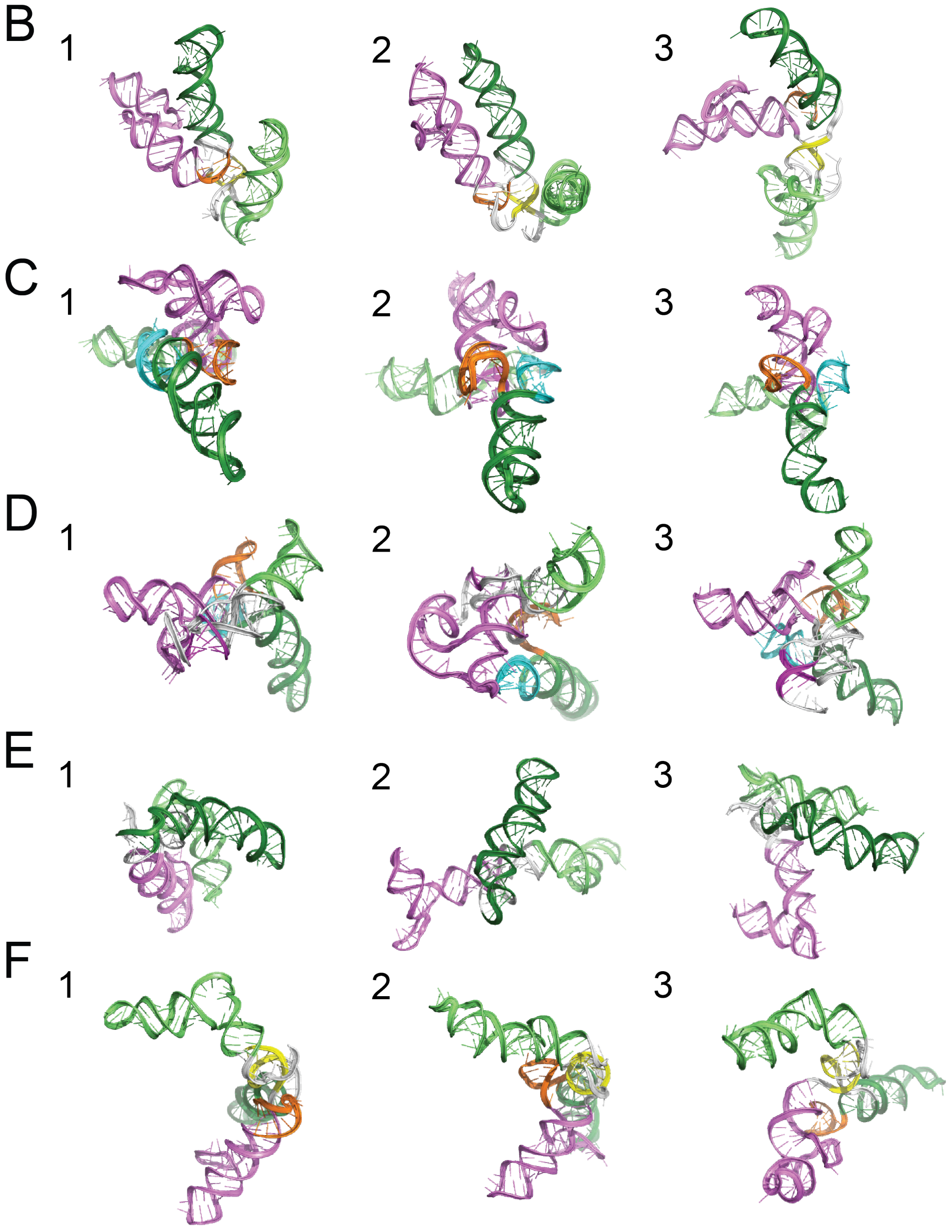


**Supplementary Figure 4.** *Models for the extended frameshift stimulation element with different secondary structures*. A) Secondary structure for the extended FSE as predicted from RNAstructure guided by the SHAPE data collected Zhang, et al.[1] for this extended FSE construct. Top 3 clusters are depicted for an extended segment including the frameshift stimulating element using secondary structures predicted by RNAstructure guided by chemical reactivity data from B) Zhang, et al.[1] C) Manfredonia, et al.[2] D) Huston, et al.[3] E) Lan, et al.[4] and F) Iserman, et al.[5]

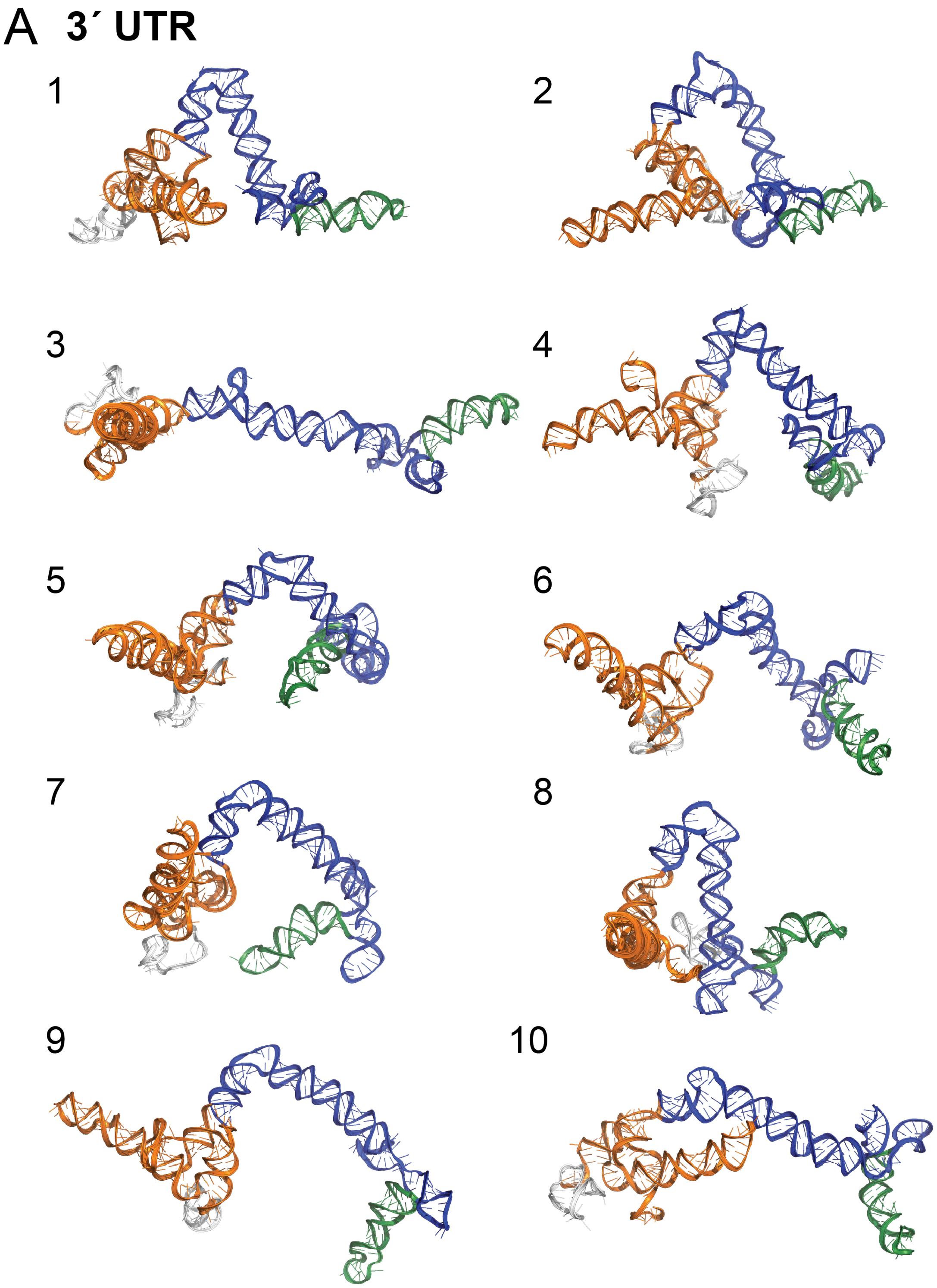

A 3´ UTR
1
2
3
4
5
6
7
8
9
10

**B Extended 3´ UTR Pseudoknot**

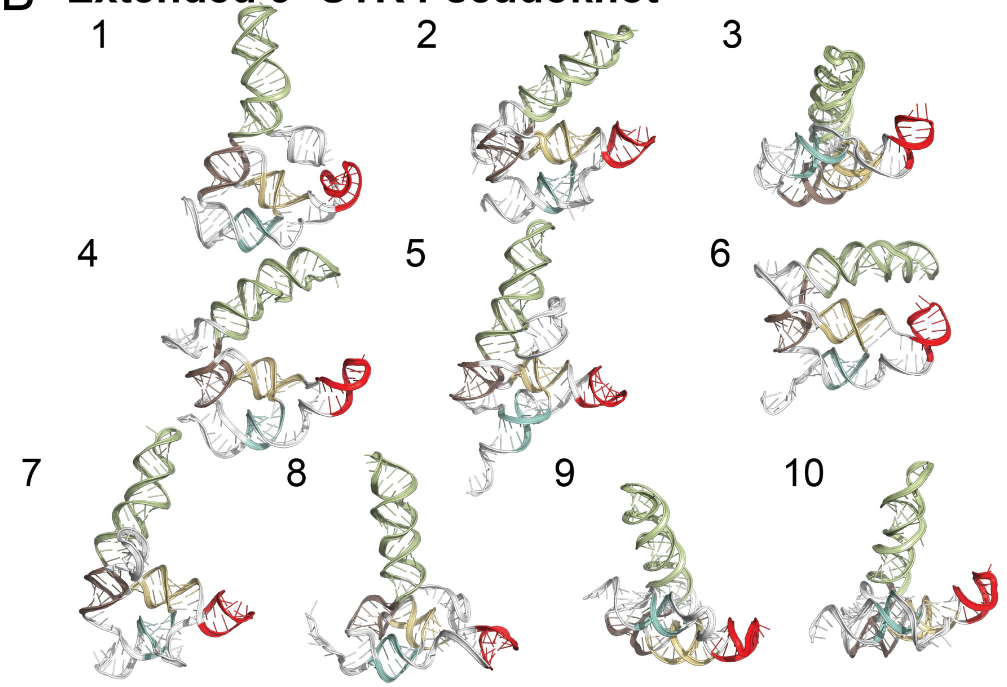


**C 3´ UTR Extended BSL**

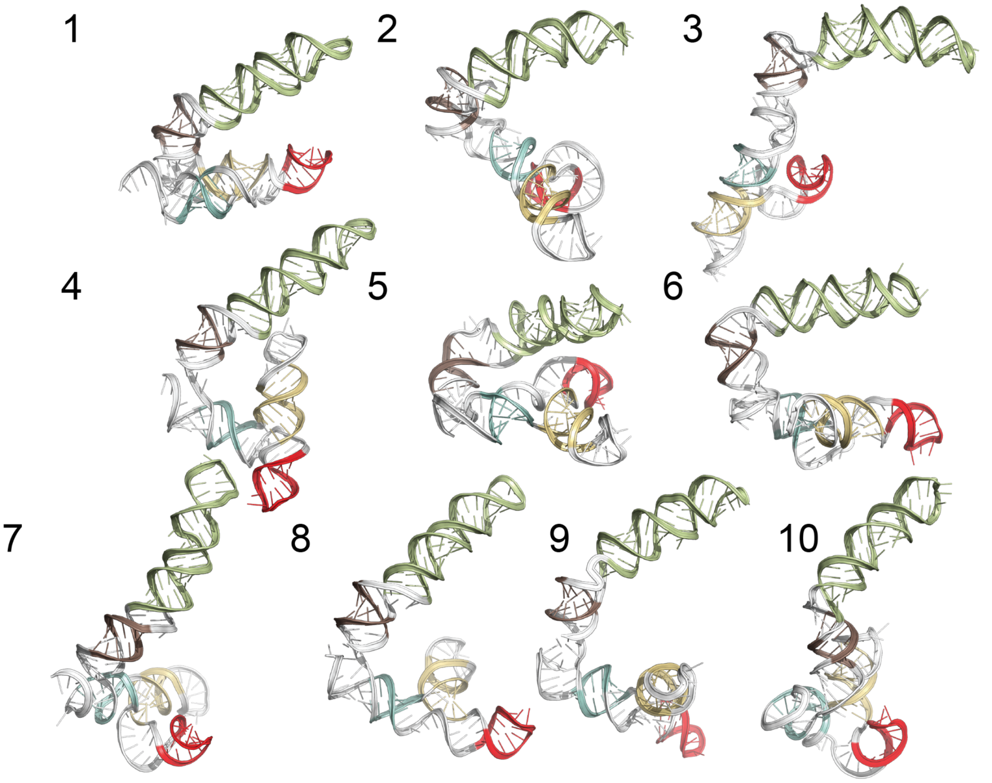

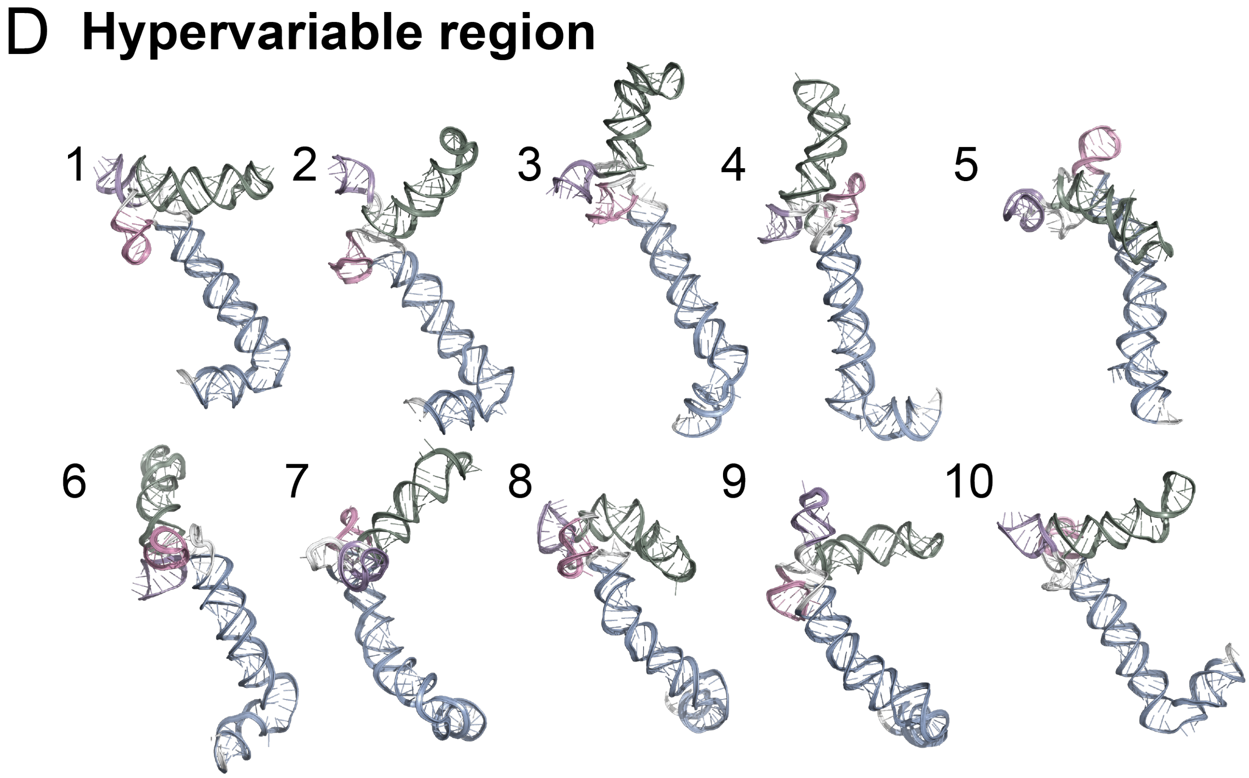


**Supplementary Figure 5.** *Models for full 3′ UTR and segments of the 3′ UTR*. Top 10 clusters (all single-occupancy) are depicted for the A) complete 3′ UTR, B) 3′ UTR pseudoknot, C) 3′ UTR extended BSL, and B) hypervariable region,. Structures are colored analogously to the 3′ UTR secondary structure in Fig. 4.

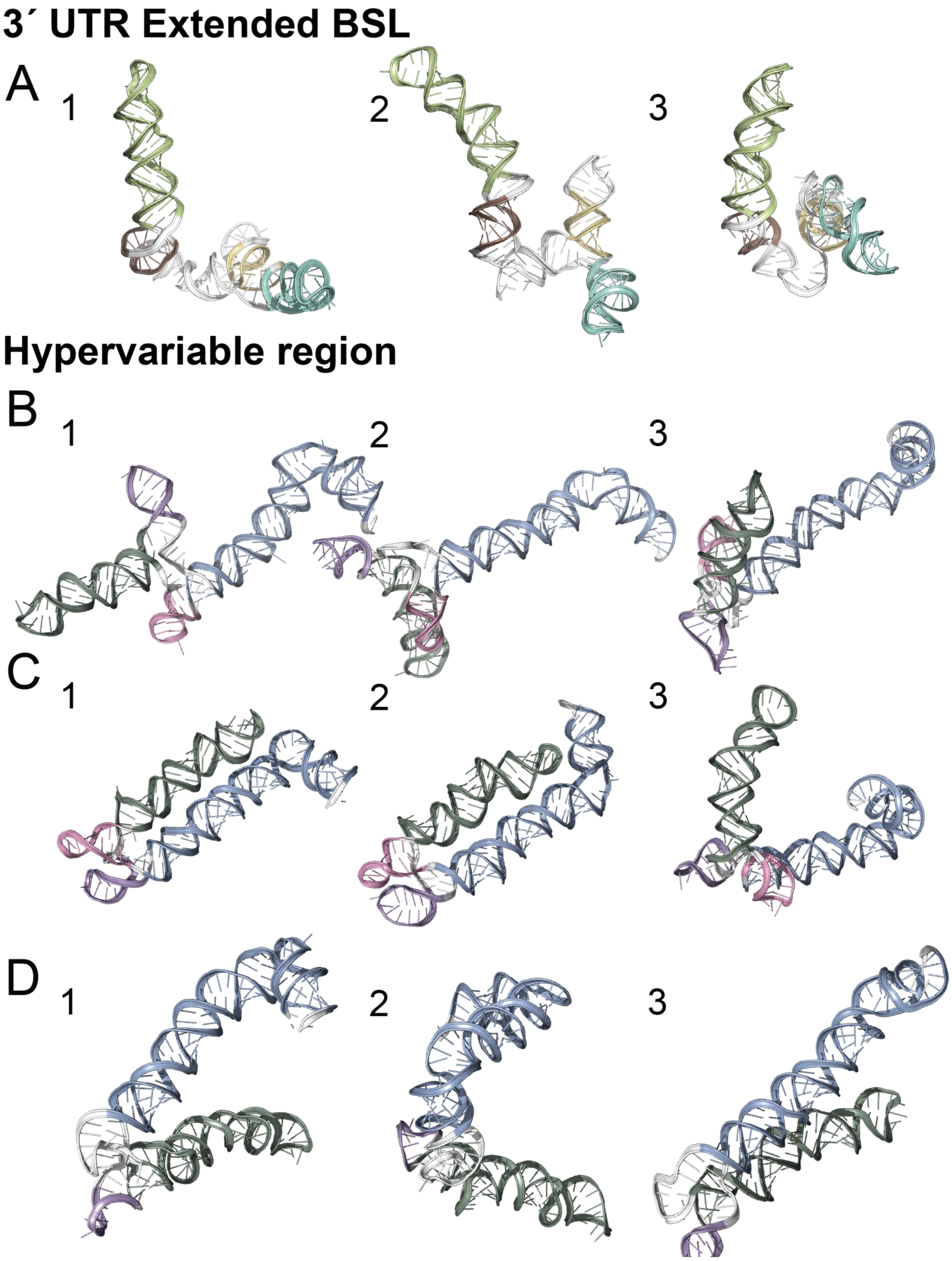


**Supplementary Figure 6.** *Models for segments of the 3′ UTR with alternate secondary structures.* A) Top 3 clusters are depicted for the 3′ UTR pseudoknot / extended BSL region with secondary structure as predicted from RNAstructure guided by chemical reactivity data from Huston, et al.[3] Top 3 clusters are depicted for the hypervariable region with structures predicted from RNAstructure guided by chemical reactivity data from B) this study (see Methods), C) Manfredonia, et al.[2], and D) Huston, et al.[3] Coloring of stems matches Fig. 4.

**HOMOLOGY**

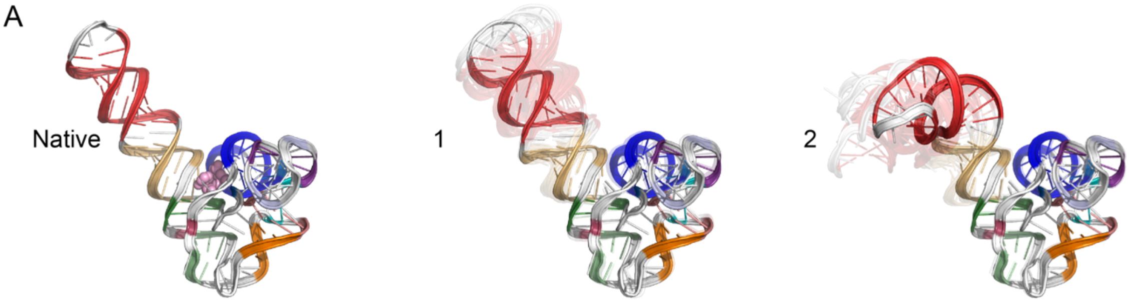


SAM-I riboswitch (RNA-Puzzle 4)

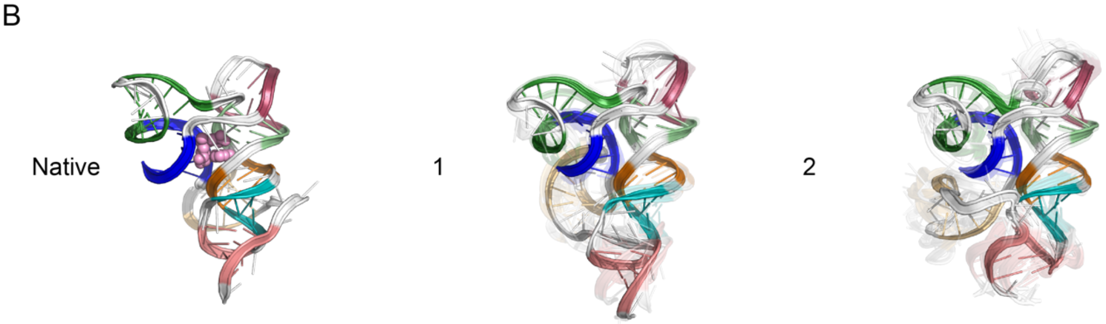


SAM-I/IV riboswitch (RNA-Puzzle 8)

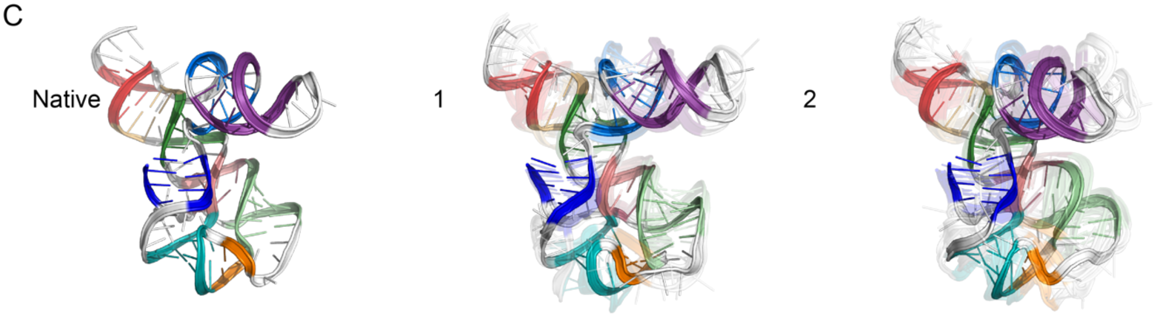


SAM-IV riboswitch (RNA-Puzzle Unknown RFam 15)

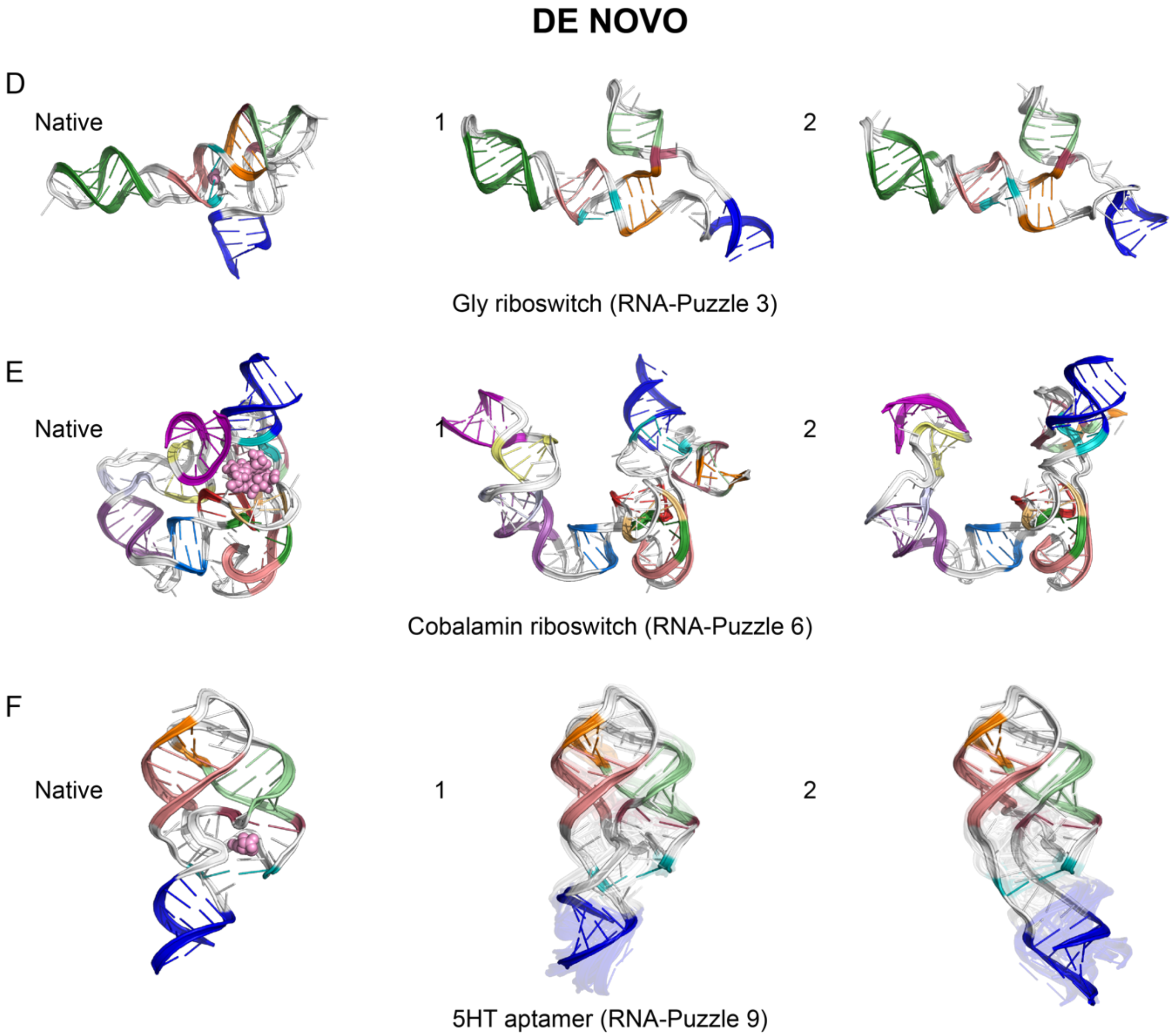
DE NOVO
D
Native
1
2
Gly riboswitch (RNA-Puzzle 3)
E
Native
1
2
Cobalamin riboswitch (RNA-Puzzle 6)
F
Native
1
2
5HT aptamer (RNA-Puzzle 9)

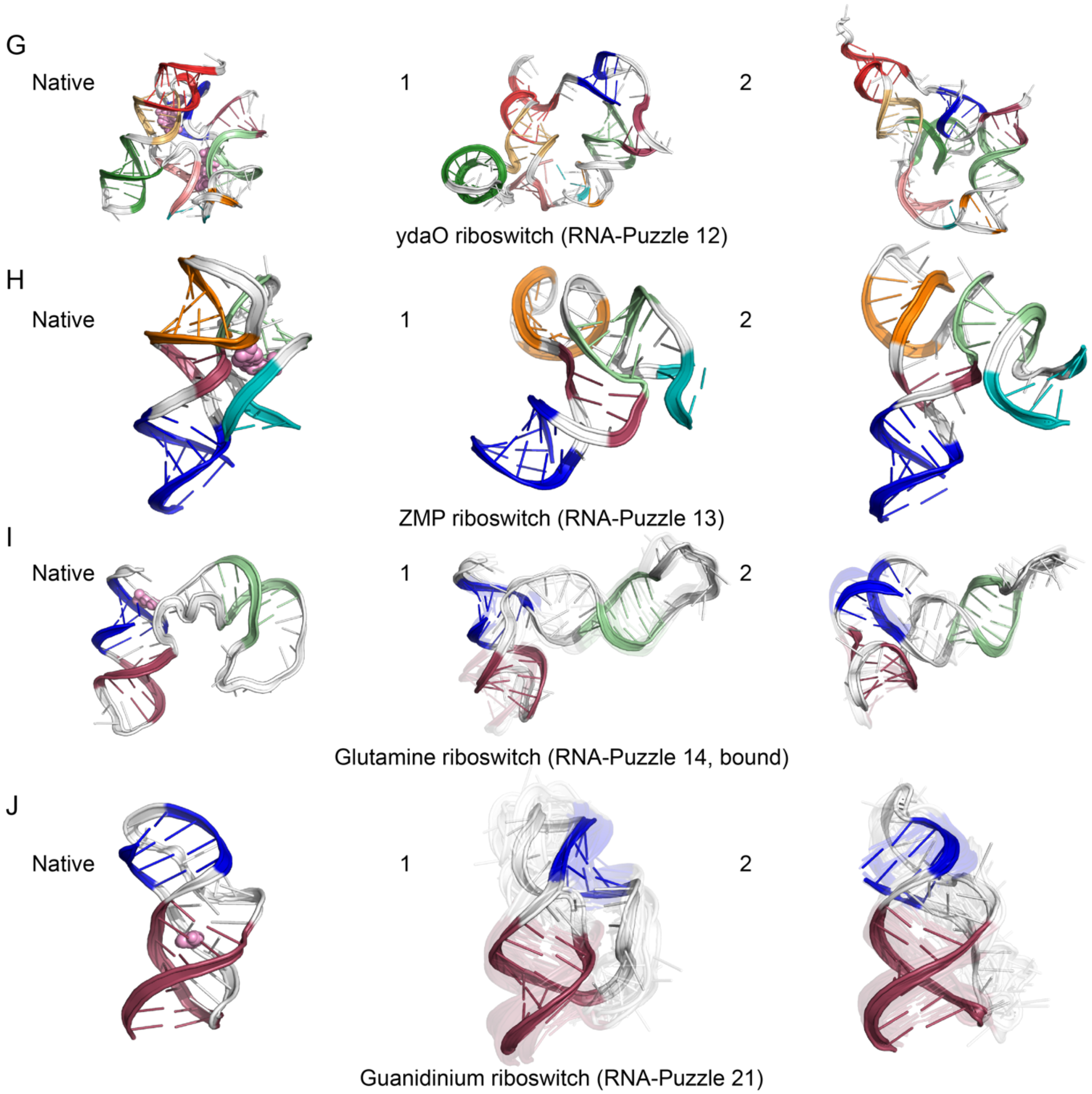


**Supplementary Figure 7.** Top two clusters for the following small-molecule binding RNA riboswitches and aptamers, along with their native structure (left in each panel, with ligand in pink spheres), beginning with three template-guided modeling challenges: A) SAM-I riboswitch, B) SAM-I/IV riboswitch, C) SAM-IV riboswitch, D) glycine riboswitch, E) cobalamin riboswitch, F) 5HT riboswitch, G) cyclic diGMP ydaO riboswitch , H) ZMP riboswitch, I) glutamine riboswitch, J) guanidinium riboswitch. The top-scoring cluster member in each case is depicted with solid colors, and the top cluster members (up to 10) are depicted as transparent structures.

**Table S1.** Secondary structures for SARS-CoV-2 genome elements.

| Region | Positions | Dataset[a] | ShapeKnots vs Fold[b] | Used for 3D models | Secondary structure |
|---|---|---|---|---|---|
| 5'UTR extended | 1-480 | Literature[6] | -- | Yes | ......(((((.(((((....)))))..)))))...........(((((.....))))).((((.......)))).<br>.......((((((((.((.((((.(((.....))).)))))).))))))))......................(((<br>((((((((..(((((...(((.(((((((((((.((((((.(((((......)))))..))))))).....)<br>))((((((((.((......))))))))(((....))))))))))))))).))))).))))...))))))).....<br>...((((((.......((..((((((...))))))..)))))))).....(((((.((((((((((((.....<br>)))).)))))..))))).)))))...........((((((((.((.((((......(((((((....(((.....<br>))))))))))....)))).)).))).))))).. |
| 5'UTR extended | 1-480 | Manfredonia, et al.[2] | Fold | No | ......(((((.(((((....)))))..)))))...........(((((.....))))).((((.......)))).<br>.......((((((((.((.((((.(((.....))).)))))).))))))))......................(((<br>((((((((..(((((...(((.(((((((((((..((((((..((((......))))...)))))))......))<br>)(((((((.((......))))))))(((....))))))))))))))).))))).))))...))))))).....<br>..((((((.......((..((((((...))))))..))))))))...(((((((.((((((((((((.....)<br>))).)))..))))))).)))))..))........((((((((.((.((((..((..(((((((....(((.....)<br>))))))))))..)))))).)).)))).)))).. |
| 5'UTR extended | 1-480 | Huston, et al.[3] | Fold | Yes: SL8 | .((((.(((((.(((((....)))))..)))))...........(((((.....))))).((((.......))))<br>.)))).((((((((((.((.((((.(((.....))).)))))).)))))))).)).........((........))<br>((((((((((..(((((...(((.(((((((((((..((((((.(((((......)))))..)))))).......<br>)))((((((((.((......))))))))(((....))))))))))))))).))))).))))...)))))).....<br>....((((((...........((((((...))))))....))))))......(((((.((((((((((((.....<br>)))).)))))..))))).)))))...........(((((((((....((((..((..((((((....((((.....<br>))))))))))))..))))))....)))).)))).. |
| 5'UTR extended | 1-480 | Iserman, et al.[5] | Fold | No | ......(((((.(((((....)))))..)))))...........(((((.....))))).((((.......)))).<br>.......((((((((.((.((((.(((.....))).)))))).))))))))......................(((<br>((((((((..(((((...(((.(((((((((((..((((((.(((((......)))))..)))))))......))<br>)(((((((.((......))))))))(((....))))))))))))))).))))).))))...))))))).....<br>..((((((.......((..((((((...))))))..)))))))).....(((((.((((((((((((.....)<br>))).))))..))))).)))))...........((((((((.((.((((..((..(((((((....(((((...))<br>))))))))))..)))))).)).))).))))).. |
| 5'UTR extended | 1-480 | Sun et al.[7], *in vivo* | Fold | No | .((((.(((((.(((((....)))))..)))))...........(((((.....))))).((((.......))))<br>......((((((((((.((.((((.(((.....))).)))))).)))))))).)).((((.....)))).....<br>((((((((((..(((((...(((.(((((((((((..((((((.(((((......)))))..))))))....<br>..)))((((((((.((......))))))))(((....))))))))))))))).))))).))))...))))))).<br>....(.((((((.......((..((((((...))))))..))))))))...).(((((.((((((((((((..<br>...)))).))))..))))).)))))........)))).((((((((.((.((((..((..(((((((....(((((<br>...))))))))))))..)))))).)).)))).)))).. |
| 5'UTR extended | 1-480 | Sun et al.[7], *in vitro* | Fold | Yes: SL8 | .((((.(((((.(((((....)))))..)))))...........(((((.....))))).((((.......))))<br>........((((((((.((.((((.(((.....))).)))))).))))))))((((((((.....)))))).))<br>((((((((((..(((((...(((.(((((((((((..((((((.(((((......)))))..))))))....<br>..)))((((((((.((......))))))))(((....))))))))))))))).))))).))))...))))))).<br>......((((((.......((..((((((...))))))..))))))))...(((((((.((((((((((((..<br>...)))).))))..))))).)))))..))...)))).((((((((.((.((((..((..(((((((....((((<br>...))))))))))))..)))))).)).))).))))).. |
| 5'UTR SL1-4 | 1-149 | This study | Fold | Yes: extended SL4 | ......(((((.(((((....)))))..)))))...........(((((.....))))).((((.......)))).<br>.......((((((((.((.((((.(((.....))).)))))).))))))))((((((((.....)))))).)) |
| 5'UTR SL2-6 | 45-344 | This study | Fold | No | (((((.....)))))......................((((((((.((.((((.(((.....))).)))))).)<br>)))))))......................((((((((((..(((((...(((.(((((((.(((..((((((.(<br>((((......)))))..))))))......)))((((((((.((......))))))))(((....))).))))))<br>)))).))))).))))...)))))))).......((((((.......((..((((((...))))))..))))))))<br>. |
| RC_TRS | 149-1 | This study | Fold | Yes | ..((((((((.....)))))))).(((((((..(.....)..)))))))(((.((((((((((.((.((((((...<br>((((.......)))).(((((.....))))).)))))).))))))).))))).)))((((((....))))))..<br>. |
| plusFSE | 13349-13546 | Manfredonia, et al.[2] | ShapeKnots | Yes | .((((((......)))).)((((((...(((.(((((.....)))).).)))...))))))(((((....)))))<br>((((((((((.(((.((<br>((....)))).)))...............))))))))))[[[[[[...((((((([]]]]]..((((((((((((<br>....)).))).))))) ))))).)))))).... |
| plusFSE | 13349-13546 | Iserman, et al.[5] | Fold | Yes | .......((((((.....((((((...(((.(((((.....)))))..)))...))))))(((((....)))))(<br>(((((((((.(((.(((<br>(....)))).)))...............))))))))))..)))))).(((((..........((((((((((...<br>......))).)))))))) ...)))))).... |
| plusFSE | 13349-13546 | Lan, et al.[4] | Fold | Yes | .......(((.....(((((((((...(((.(((((.....)))))..)))...))))))).)))))).....(.(<br>(((((((((.(((.(((<br>(....)))).)))...............)))))))))).).......((((..........((((((((((((..<br>..)).))).))))))))) ...)))).....<br> |

| | | | | | |
|---|---|---|---|---|---|
| plusFSE | 13349-13546 | Huston, et al.[3] | ShapeKnots | Yes | .(((((.......)))))((((((...(((.(((((.....)))))..)))...))))))(((((....)))))...(((((((.(((.((((....)))).)))...............)))))))((((((((...[[[[[..)))))))))((((((((((.........))).)))))))....]]]]].... |
| plusFSE | 13349-13546 | Sun et al.[7], *in vivo* | Fold | Yes | ((((...((((((.....((((((((.(((.((((((...))))))..))))).))))))(((((....)))))((((((((((.(((.((((....)))).))).....(((....))))))))))))..)))))).)))).(((((.....((((((((((((....)).))).))))))))..))))).... |
| plusFSE | 13349-13546 | Sun et al.[7], *in vitro* | ShapeKnots | Yes | (((.....((((((.....((((((...(((.((((((...))))))..)))...))))))(((((....)))))((((((((((.(((.((((....)))).))).....(((....))))))))))))..))))))).[[[[[[[.....)))((((((((((((((....)).))).))))))))]].]]]]].... |
| plusFSE | 13349-13546 | Zhang et al.[1] | Fold | Yes | .....((((.....((((((((...(((..(.((.....)).)...)))...)))))).)).(.....).((.((((((((((..((.((((....)))).)).(.....).........))))))))).)).)))).((((((..........(((((((((((.... .....))).))))))))...))))).... |
| FSE | 13459-13546 | Literature[8] | -- | Yes | ...............((((((((((...[[[[[[[))))))))))((((((((((.........))).))))))..]].]]]]].... |
| FSE | 13459-13546 | Literatre[6] | -- | Yes | ..............(((((((((((...[[[[[[[)))))))))))((((((((.........))).)))))...]].]]]]].... |
| Pkfull | 29543–29665; 29846–29876 | Literature[6] (pseudoknot) | -- | Yes | ...............(((((((((((..(((...((......))...))).))))))))))..((((((((.......{{{{{{.[[[[[[[[[.))))))))...]]]]]]]]]...((((......))))...........}}}}}}......... |
| Pkfull | 29543–29665; 29846–29876 | Literature[6] (BSL ext) | -- | Yes | .....((((((.....(((((((((((..(((...((......))...))).))))))))))..))))))........{{{{{{.(((((((((...........)))))))))...((((......))))...........}}}}}}......... |
| Pkfull | 29543–29665; 29846–29876 | Manfredonia, et al.[2] | Fold | No | .....((((((..((.(((((((((((..(((...((......))...))).)))))))))))))))))))........((((((.(((((((((.((....))..))))))))).((((((......)))).)).........))))))......... |
| Pkfull | 29543–29665; 29846–29876 | Manfredonia, et al.[2] | ShapeKnots | No | .....((((((..((.(((((((((((..(((...((......))...))).)))))))))))))))))))........((((((.(((((((((.((....))..))))))))).((((((......)))).)).........))))))......... |
| Pkfull | 29543–29665; 29846–29876 | Huston, et al.[3] | Fold | No | .....((((((..((.(((((((((((..(((...((......))...))).)))))))))))))))))))).((........))..(((((((((...........))))))))).((((((.(((.((....)).))))))).))............ |
| Pkfull | 29543–29665; 29846–29876 | Huston, et al.[3] | ShapeKnots | Yes | .....((((((..((.(((((((((((..(((...((......))...))).)))))))))))))))))))...............(((((((((...........))))))))).((((((.(((..........))))))).))............ |
| Pkfull | 29543–29665; 29846–29876 | Sun et al.[7], *in vivo* | Fold | No | .....(((((((..(.(((((((((((..(((...((......))...))).)))))))))))))))))))........((((((.(((((((((.((....))..))))))))).((((((......)))).))((....))))))))......... |
| Pkfull | 29543–29665; 29846–29876 | Sun et al.[7], *in vivo* | ShapeKnots | No | .....(((((((..(.(((((((((((..(((...((......))...))).)))))))))))))))))))........((((((.(((((((((.((....))..))))))))).((((((......)))).))((....))))))))......... |
| Pkfull | 29543–29665; 29846–29876 | Sun et al.[7], *in vitro* | Fold | No | .....(((((((..(.(((((((((((..(((...((......))...))).)))))))))))))))))))........((((((.(((((((((.((....))..))))))))).((((((......)))).))((....))))))))......... |
| Pkfull | 29543–29665; 29846–29876 | Sun et al.[7], *in vivo* | ShapeKnots | No | .....(((((((..(.(((((((((((..(((...((......))...))).)))))))))))))))))))........((((((.(((((((((.((....))..))))))))).((((((......)))).))((....))))))))......... |
| HVR | 29659–29852 | Literature[6] | -- | Yes | ..((((..((((((((((..((..(((.(((.....((((((((.((.(((....)))))....((((((((....((..((((.....))..))...))...))))).)))).((((.........))))..........))))))))).....))).)))..))...)))))......)))))..)))).. |
| HVR | 29659–29852 | Manfredonia, et al.[2] | Fold | Yes | ..((((..((((((((((..((..(((.(((.....((((((((.((.(((....)))))).((.(((((((((....((.((.((.(.......))).))))...))))).))))...))......((.((.....)).))..))))))))))......))).)))..))...))))......))))))..)))).. |
| HVR | 29659–29852 | Huston, et al.[3] | Fold | Yes | ..((((..((((((((((..((..(((.........((((((((............(((..((.(((((((((....((.((...((......))..))))...))))).))))...))....))).((.....)).....))))))))).........)))..))...))))))......))))..)))).. |
| HVR | 29659–29852 | This study | Fold | Yes | ..((((..((((((((((..((..(((.(((.....((((((((.((.(((....)))))....((((((((....((.((.(((........))).))))...))))).)))).((((.........))))..........))))))))).....))).)))..))...))))......))))))..)))).. |

| | | | | | |
|---|---|---|---|---|---|
| HVR | 29659–29852 | Sun et al.[7], *in vivo* | Fold | No | ..((((..((((((((((..((..(((.(((.....((((((((.((.(((....))))).((.((((((((<br>(....(.(((.(((.......))).))))...))))).))))...)).....((.((.....)).))..)))))))<br>)).....))).)))..))...))))......))))))..)))).. |
| HVR | 29659–29852 | Sun et al.[7], *in vitro* | Fold | No | ..((((..((((((((((..((..(((.(((.....((((((((.((.(((....))))).((((((((((((<br>(....(.(((.(((.......))).))))...))))).)))).((((........))))..))).....))))))<br>))).....))).)))..))...))))......))))))..)))).. |
| 3'UTR | 29543–29903 | Literature[6] | -- | Yes | ................((((((((((((..(((...((......))...))).)))))))))))..(((((((....<br>...{{{{{{.[[[[[[[[[.))))))))...]]]]]]]]]...((((..((((((((((..((..(((.(((.....(<br>((((((((.((.(((....)))))....(((((((((....((..((((.....))..))...))...)))))).)<br>))).((((........))))..........)))))))))).....))).)))..))...)))))......)))))..<br>))))...........}}}}}}...................................... |
| s2m | 29724-29773 | Wacker, et al.[9] | -- | Yes | (((((......(((.((.........)).)))....))))) |
| s2m | 29724-29773 | Manfredonia, et al.[2] | Fold | Yes | (((((....((.((.((.(......))).))))...))))) |
| s2m | 29724-29773 | This study | Fold | Yes | (((((....((.((.(((.......))).))))...))))) |
| s2m | 29724-29773 | Huston, et al.[3] | Fold | Yes | (((((....((.((...((......))..))))...))))) |

[a]Chemical probing dataset used to model secondary structure. \`Literature\` indicates that structure was obtained from consensus structures reported in prior research.
[b]RNAstructure predictions guided by chemical reactivity data were run with Fold (no pseudoknots allowed) or ShapeKnots (pseudoknots predicted).

**Table S2.** FARFAR2-SARS-CoV-2 models, extended data.

| System/cluster | Length | Models generated | Model convergence (Å)[a] | Predicted Minimum RMSD (Å)[b] | E-gap to lowest energy model (REU)[c] | Cluster occupancy[d] |
|---|---|---|---|---|---|---|
| **Extended 5′ UTR (1–480)** | | | | | | |
| **5′ UTR/1** | 480 | 66011 | 50.9 | 44.92 ± 6.13 | 0 | 1 |
| 5′ UTR/2 | | | | | 5.15 | 1 |
| 5′ UTR/3 | | | | | 12.45 | 1 |
| 5′ UTR/4 | | | | | 12.66 | 1 |
| 5′ UTR/5 | | | | | 12.98 | 1 |
| 5′ UTR/6 | | | | | 13.07 | 1 |
| 5′ UTR/7 | | | | | 13.92 | 1 |
| 5′ UTR/8 | | | | | 14.9 | 1 |
| 5′ UTR/9 | | | | | 15.43 | 1 |
| 5′ UTR/10 | | | | | 21.01 | 1 |
| **5′ UTR stem-loop 1 (7–33)** | | | | | | |
| SL1/1 | 27 | 200000 | 1.83 | 5.17 ± 0.52 | 0 | 6 |
| SL1/2 | | | | | 0.07 | 16 |
| SL1/3 | | | | | 0.26 | 45 |
| SL1/4 | | | | | 0.91 | 13 |
| SL1/5 | | | | | 0.97 | 6 |
| SL1/6 | | | | | 1.02 | 18 |
| SL1/7 | | | | | 1.04 | 9 |
| SL1/8 | | | | | 1.27 | 14 |
| SL1/9 | | | | | 1.55 | 4 |
| SL1/10 | | | | | 1.57 | 12 |
| **5′ UTR stem-loop 2 (45–59)** | | | | | | |
| **SL2/1** | 15 | 200000 | 2.39 | 5.63 ± 0.74 | 0 | 52 |
| SL2/2 | | | | | 0.73 | 44 |
| SL2/3 | | | | | 0.95 | 78 |
| SL2/4 | | | | | 0.98 | 56 |

| | | | | | | |
|---|---|---|---|---|---|---|
| SL2/5 | | | | | 1.08 | 15 |
| SL2/6 | | | | | 1.13 | 207 |
| SL2/7 | | | | | 1.15 | 26 |
| SL2/8 | | | | | 1.38 | 29 |
| SL2/9 | | | | | 1.4 | 13 |
| SL2/10 | | | | | 1.48 | 41 |
| SL2/Homology | | | | | 20.2 | 27 |
| **5′ UTR stem-loop 3 (61–75)** | | | | | | |
| **SL3/1** | 15 | 200000 | 2.58 | 5.78 ± 0.70 | 0 | 7 |
| SL3/2 | | | | | 0.1 | 45 |
| SL3/3 | | | | | 0.61 | 10 |
| SL3/4 | | | | | 0.78 | 69 |
| SL3/5 | | | | | 1.31 | 70 |
| SL3/6 | | | | | 1.55 | 6 |
| SL3/7 | | | | | 1.89 | 84 |
| SL3/8 | | | | | 1.93 | 49 |
| SL3/9 | | | | | 2.1 | 35 |
| SL3/10 | | | | | 2.23 | 18 |
| **5′ UTR stem-loop 4 (84–127)** | | | | | | |
| **SL4/1** | 44 | 2018457 | 1.82 | 5.16 ± 0.49 | 0 | 5 |
| SL4/2 | | | | | 0.05 | 1 |
| SL4/3 | | | | | 0.91 | 4 |
| SL4/4 | | | | | 0.95 | 4 |
| SL4/5 | | | | | 1.03 | 5 |
| SL4/6 | | | | | 1.16 | 11 |
| SL4/7 | | | | | 1.43 | 6 |
| SL4/8 | | | | | 1.74 | 2 |
| SL4/9 | | | | | 1.91 | 3 |
| SL4/10 | | | | | 1.92 | 1 |
| **5′ UTR stem-loop 5 (148–295)** | | | | | | |

| | | | | | | |
|---|---|---|---|---|---|---|
| **SL5/1** | 148 | 2392320 | 18.99 | 19.07 ± 6.15 | 0 | 1 |
| SL5/2 | | | | | 1.39 | 2 |
| SL5/3 | | | | | 1.46 | 1 |
| SL5/4 | | | | | 2.31 | 1 |
| SL5/5 | | | | | 2.32 | 1 |
| SL5/6 | | | | | 3.1 | 1 |
| SL5/7 | | | | | 3.35 | 1 |
| SL5/8 | | | | | 3.87 | 1 |
| SL5/9 | | | | | 4.16 | 1 |
| SL5/10 | | | | | 4.23 | 1 |
| **5′ UTR stem-loop 5/6 (148–343)** | | | | | | |
| **SL56/1** | 196 | 2020963 | 25.91 | 24.68 ± 6.27 | 0 | 1 |
| SL56/2 | | | | | 4.42 | 1 |
| SL56/3 | | | | | 4.89 | 2 |
| SL56/4 | | | | | 5.69 | 1 |
| SL56/5 | | | | | 6.75 | 1 |
| SL56/6 | | | | | 8.16 | 1 |
| SL56/7 | | | | | 8.21 | 1 |
| SL56/8 | | | | | 8.47 | 1 |
| SL56/9 | | | | | 8.52 | 1 |
| SL56/10 | | | | | 8.92 | 1 |
| **5′ UTR stem-loop 6 (302–343)** | | | | | | |
| **SL6/1** | 42 | 200000 | 8.92 | 10.92 ± 2.21 | 0 | 162 |
| SL6/2 | | | | | 4.21 | 537 |
| SL6/3 | | | | | 4.97 | 27 |
| SL6/4 | | | | | 5.32 | 12 |
| SL6/5 | | | | | 5.37 | 119 |
| SL6/6 | | | | | 5.38 | 193 |
| SL6/7 | | | | | 6.03 | 15 |
| SL6/8 | | | | | 6.42 | 47 |
| SL6/9 | | | | | 6.64 | 26 |

| | | | | | | |
|---|---|---|---|---|---|---|
| SL6/10 | | | | | 6.86 | 148 |
| **5′ UTR stem-loop 7 (349–394)** | | | | | | |
| **SL7/1** | 27 | 200000 | 7.39 | 9.67 ± 1.90 | 0 | 256 |
| SL7/2 | | | | | 1.35 | 292 |
| SL7/3 | | | | | 12.24 | 726 |
| SL7/4 | | | | | 14.6 | 15 |
| SL7/5 | | | | | 17.5 | 46 |
| SL7/6 | | | | | 20.22 | 2449 |
| SL7/7 | | | | | 23.11 | 9 |
| SL7/8 | | | | | 23.61 | 20 |
| SL7/9 | | | | | 23.88 | 43 |
| SL7/10 | | | | | 25.41 | 331 |
| **5′ UTR stem-loop 8 (407-478)** | | | | | | |
| **SL8/1** | 72 | 4055322 | 6.86 | 9.24 ± 1.47 | 0 | 3 |
| SL8/2 | | | | | 2.51 | 7 |
| SL8/3 | | | | | 3.11 | 18 |
| SL8/4 | | | | | 6.27 | 15 |
| SL8/5 | | | | | 6.96 | 47 |
| SL8/6 | | | | | 7.55 | 27 |
| SL8/7 | | | | | 7.7 | 21 |
| SL8/8 | | | | | 8.06 | 26 |
| SL8/9 | | | | | 8.69 | 17 |
| SL8/10 | | | | | 10.55 | 32 |
| **5′ UTR reverse complement stem-loops1-4 (149 - 1)** | | | | | | |
| **RC-SL1-4/1** | 149 | 2031710 | 19.61 | 19.57 ± 4.37 | 0 | 1 |
| RC-SL1-4/2 | | | | | 1.5 | 1 |
| RC-SL1-4/3 | | | | | 3.83 | 1 |
| RC-SL1-4/4 | | | | | 4.41 | 1 |
| RC-SL1-4/5 | | | | | 4.5 | 1 |
| RC-SL1-4/6 | | | | | 4.78 | 1 |

| | | | | | | |
|---|---|---|---|---|---|---|
| RC-SL1-4/7 | | | | | 4.8 | 1 |
| RC-SL1-4/8 | | | | | 6.01 | 1 |
| RC-SL1-4/9 | | | | | 7.78 | 1 |
| RC-SL1-4/10 | | | | | 9.21 | 1 |
| **Frameshift stimulating element (13459–13546)** | | | | | | |
| **FSE/1** | 88 | 390722 | 14.45 | 15.39 ± 3.09 | 0 | 5 |
| FSE/2 | | | | | 3.23 | 1 |
| FSE/3 | | | | | 4.44 | 2 |
| FSE/4 | | | | | 5.44 | 3 |
| FSE/5 | | | | | 6.22 | 2 |
| FSE/6 | | | | | 7.47 | 2 |
| FSE/7 | | | | | 7.78 | 1 |
| FSE/8 | | | | | 7.81 | 4 |
| FSE/9 | | | | | 8.06 | 11 |
| FSE/10 | | | | | 8.17 | 8 |
| **Suspected frameshift stimulating element dimer (13459–13546 x 2)** | | | | | | |
| **FSE Dimer/1** | 176 | 23066 | 21.99 | 21.50 ± 4.08 | 0 | 1 |
| FSE Dimer/2 | | | | | 6.37 | 1 |
| FSE Dimer/3 | | | | | 6.79 | 1 |
| FSE Dimer/4 | | | | | 10.02 | 1 |
| FSE Dimer/5 | | | | | 10.11 | 1 |
| FSE Dimer/6 | | | | | 10.22 | 1 |
| FSE Dimer/7 | | | | | 10.31 | 1 |
| FSE Dimer/8 | | | | | 11.55 | 1 |
| FSE Dimer/9 | | | | | 12.06 | 1 |
| FSE Dimer/10 | | | | | 12.45 | 1 |
| **3′ UTR beginning with bulged hairpin (29511-29871)** | | | | | | |
| **3′ UTR/1** | 361 | 11430 | 39.71 | 35.85 ± 5.51 | 0 | 1 |
| 3′ UTR/2 | | | | | 4.07 | 1 |
| 3′ UTR/3 | | | | | 26.33 | 1 |

| | | | | | | |
|---|---|---|---|---|---|---|
| 3′ UTR/4 | | | | | 32.67 | 1 |
| 3′ UTR/5 | | | | | 42.1 | 1 |
| 3′ UTR/6 | | | | | 43.14 | 1 |
| 3′ UTR/7 | | | | | 46.96 | 1 |
| 3′ UTR/8 | | | | | 47.55 | 1 |
| 3′ UTR/9 | | | | | 48.04 | 1 |
| 3′ UTR/10 | | | | | 48.28 | 1 |
| **3´ UTR hypervariable region (29659-29852)** | | | | | | |
| **HVR/1** | 194 | 28029 | 25.38 | 24.25 ± 4.18 | 0 | 1 |
| HVR/2 | | | | | 2.49 | 1 |
| HVR/3 | | | | | 3.6 | 1 |
| HVR/4 | | | | | 4.16 | 1 |
| HVR/5 | | | | | 5.81 | 1 |
| HVR/6 | | | | | 6.79 | 1 |
| HVR/7 | | | | | 7.63 | 1 |
| HVR/8 | | | | | 7.77 | 1 |
| HVR/9 | | | | | 9.06 | 1 |
| HVR/10 | | | | | 11.08 | 1 |
| **3´ UTR pseudoknot (29543–29665; 29846–29876)** | | | | | | |
| **Pseudoknot/1** | 158 | 1017205 | 21.93 | 21.45 ± 5.52 | 0 | 1 |
| Pseudoknot/2 | | | | | 1.42 | 1 |
| Pseudoknot/3 | | | | | 2.87 | 1 |
| Pseudoknot/4 | | | | | 3.94 | 1 |
| Pseudoknot/5 | | | | | 3.95 | 1 |
| Pseudoknot/6 | | | | | 4.06 | 1 |
| Pseudoknot/7 | | | | | 4.27 | 1 |
| Pseudoknot/8 | | | | | 4.97 | 1 |
| Pseudoknot/9 | | | | | 5.37 | 1 |
| Pseudoknot/10 | | | | | 6.15 | 1 |
| **3´ UTR pseudoknot fragment consisting of the pseudoknot (PK), P2, and P5 (29606–29665; 29842–29876)** | | | | | | |

| | | | | | | |
|---|---|---|---|---|---|---|
| **PK-P2-P5/1** | 95 | 1017205 | 10.24 | 11.99 ± 2.05 | 0 | 1 |
| PK-P2-P5/2 | | | | | 0.64 | 2 |
| PK-P2-P5/3 | | | | | 0.64 | 1 |
| PK-P2-P5/4 | | | | | 0.69 | 2 |
| PK-P2-P5/5 | | | | | 0.98 | 3 |
| PK-P2-P5/6 | | | | | 1.02 | 5 |
| PK-P2-P5/7 | | | | | 1.51 | 1 |
| PK-P2-P5/8 | | | | | 1.53 | 1 |
| PK-P2-P5/9 | | | | | 2.43 | 1 |
| PK-P2-P5/10 | | | | | 2.82 | 3 |
| **3´ UTR BSL extended structure (29543–29665; 29846–29876)** | | | | | | |
| **BSLext/1** | 158 | 1012716 | 24.04 | 23.16 ± 4.29 | 0 | 1 |
| BSLext/2 | | | | | 0.48 | 1 |
| BSLext/3 | | | | | 2.45 | 1 |
| BSLext/4 | | | | | 6.22 | 1 |
| BSLext/5 | | | | | 6.34 | 1 |
| BSLext/6 | | | | | 6.38 | 1 |
| BSLext/7 | | | | | 6.45 | 1 |
| BSLext/8 | | | | | 6.56 | 1 |
| BSLext/9 | | | | | 6.76 | 1 |
| BSLext/10 | | | | | 6.84 | 1 |
| **3´ UTR stem-loop II-like motif, homology modeled from PDB ID: 1XJR[10] (29724–29773)** | | | | | | |
| **S2M/1** | 50 | 200000 | 6.95 | 9.32 ± 0.09 | 0 | 199977 |
| S2M/2 | | | | | 11.38 | 23 |
| **3´ UTR stem-loop II-like motif, secondary structure from NMR data in Wacker, et al.[9] (29724–29773)** | | | | | | |
| **S2M/1** | 50 | 500000 | 2.97 | 6.10 ± 0.75 | 0 | 1 |
| S2M/2 | | | | | 0.63 | 2 |
| S2M/3 | | | | | 0.63 | 1 |
| S2M/4 | | | | | 1.9 | 1 |
| S2M/5 | | | | | 2.16 | 1 |

| | | | | | | |
|---|---|---|---|---|---|---|
| S2M/6 | | | | | 2.6 | 1 |
| S2M/7 | | | | | 3.1 | 1 |
| S2M/8 | | | | | 3.45 | 1 |
| S2M/9 | | | | | 4.14 | 1 |
| S2M/10 | | | | | 4.29 | 1 |

[a]Mean pairwise all-heavy-atom RMSD between 10 lowest energy cluster centers.
[b]Predicted RMSD to true structure.
[c]Rosetta all-atom free energy gap of cluster's lowest energy model compared to lowest energy model discovered in run. REU = Rosetta energy units, calibrated so that 1.0 corresponds approximately to 1 $k_BT$.
[d]Number of models that appear in each cluster. Clustering was carried out on top 400 models ranked by Rosetta all-atom free energy, based on 5.0 Å threshold, except for small RNAs (SL1-4, SL6-7, s2m), where 2.0 Å threshold was applied.

**Table S3.** FARFAR2-SARS-CoV-2 models, extended data for alternate secondary structures.

| System/cluster | Length | Models generated | Model convergence (Å)[a] | Predicted minimum RMSD (Å)[b] | E-gap to lowest energy model (REU)[c] | Cluster occupancy[d] |
|---|---|---|---|---|---|---|
| **5′ UTR stem-loop 4 extended (84–146)** | | | | | | |
| **SL4ext/1** | 63 | 143897 | 7.5 | 9.76 ± 1.97 | 0 | 645 |
| SL4ext/2 | | | | | 5.28 | 377 |
| SL4ext/3 | | | | | 5.78 | 21 |
| SL4ext/4 | | | | | 8.29 | 17 |
| SL4ext/5 | | | | | 8.79 | 202 |
| SL4ext/6 | | | | | 9.07 | 560 |
| SL4ext/7 | | | | | 11.76 | 15 |
| SL4ext/8 | | | | | 11.83 | 25 |
| SL4ext/9 | | | | | 12.02 | 177 |
| SL4ext/10 | | | | | 12.58 | 742 |
| **FSE extended, secondary structure from Zhang, et al.[1] (13349-13546)** | | | | | | |
| **FSEext/1** | 198 | 993756 | 28.69 | 26.93 ± 4.46 | 0 | 1 |
| FSEext/2 | | | | | 1.49 | 1 |
| FSEext/3 | | | | | 2.28 | 1 |
| FSEext/4 | | | | | 4.83 | 1 |
| FSEext/5 | | | | | 5.91 | 1 |
| FSEext/6 | | | | | 5.98 | 1 |
| FSEext/7 | | | | | 6.69 | 1 |
| FSEext/8 | | | | | 7.11 | 1 |
| FSEext/9 | | | | | 7.88 | 1 |
| FSEext/10 | | | | | 8.05 | 1 |
| **FSE extended, secondary structure from Manfredonia, et al.[2] (13349-13546)** | | | | | | |
| **FSEext/1** | 198 | 1021344 | 24.16 | 23.26 ± 4.92 | 0 | 1 |
| FSEext/2 | | | | | 3.77 | 1 |
| FSEext/3 | | | | | 4.3 | 1 |
| FSEext/4 | | | | | 4.76 | 2 |

| | | | | | | |
|---|---|---|---|---|---|---|
| FSEext/5 | | | | | 4.87 | 1 |
| FSEext/6 | | | | | 5.54 | 1 |
| FSEext/7 | | | | | 5.73 | 1 |
| FSEext/8 | | | | | 6.24 | 1 |
| FSEext/9 | | | | | 6.7 | 1 |
| FSEext/10 | | | | | 7.39 | 1 |
| **FSE extended, secondary structure from Huston, et al.[3] (13349-13546)** | | | | | | |
| **FSEext/1** | 198 | 252971 | 20.79 | 20.53 ± 6.00 | 0 | 1 |
| FSEext/2 | | | | | 2.32 | 1 |
| FSEext/3 | | | | | 9 | 1 |
| FSEext/4 | | | | | 11.17 | 1 |
| FSEext/5 | | | | | 11.93 | 1 |
| FSEext/6 | | | | | 12.16 | 1 |
| FSEext/7 | | | | | 12.62 | 1 |
| FSEext/8 | | | | | 13.3 | 1 |
| FSEext/9 | | | | | 14 | 1 |
| FSEext/10 | | | | | 14.06 | 1 |
| **FSE extended, secondary structure from Lan, et al.[2] (13349-13546)** | | | | | | |
| **FSEext/1** | 198 | 267963 | 31.76 | 29.42 ± 4.62 | 0 | 1 |
| FSEext/2 | | | | | 5.1 | 1 |
| FSEext/3 | | | | | 5.23 | 1 |
| FSEext/4 | | | | | 8.23 | 1 |
| FSEext/5 | | | | | 8.25 | 1 |
| FSEext/6 | | | | | 9.59 | 1 |
| FSEext/7 | | | | | 10.78 | 1 |
| FSEext/8 | | | | | 11.69 | 1 |
| FSEext/9 | | | | | 12.69 | 1 |
| FSEext/10 | | | | | 13.24 | 1 |
| **FSE extended, secondary structure from Iserman, et al.[5] (13349-13546)** | | | | | | |
| **FSEext/1** | 198 | 1013658 | 28.16 | 26.50 ± 5.21 | 0 | 1 |

| | | | | | | |
|---|---|---|---|---|---|---|
| FSEext/2 | | | | | 0.27 | 1 |
| FSEext/3 | | | | | 1.78 | 1 |
| FSEext/4 | | | | | 2.02 | 1 |
| FSEext/5 | | | | | 2.09 | 1 |
| FSEext/6 | | | | | 2.94 | 1 |
| FSEext/7 | | | | | 4.73 | 1 |
| FSEext/8 | | | | | 5.06 | 1 |
| FSEext/9 | | | | | 5.52 | 1 |
| FSEext/10 | | | | | 5.89 | 1 |
| **HVR, secondary structure based on chemical reactivity from this study (29659– 29852)** | | | | | | |
| **HVRalt/1** | 194 | 1010583 | 28.17 | 26.51 ± 6.39 | 0 | 1 |
| HVRalt/2 | | | | | 5.15 | 1 |
| HVRalt/3 | | | | | 6.85 | 1 |
| HVRalt/4 | | | | | 6.97 | 1 |
| HVRalt/5 | | | | | 9.3 | 1 |
| HVRalt/6 | | | | | 10 | 1 |
| HVRalt/7 | | | | | 10.29 | 1 |
| HVRalt/8 | | | | | 11.23 | 1 |
| HVRalt/9 | | | | | 11.4 | 1 |
| HVRalt/10 | | | | | 11.44 | 1 |
| **HVR, secondary structure from Manfredonia, et al.**[2] **(29659– 29852)** | | | | | | |
| **HVRalt/1** | 194 | 1011993 | 25.11 | 24.03 ± 5.66 | 0 | 1 |
| HVRalt/2 | | | | | 3.31 | 1 |
| HVRalt/3 | | | | | 6.86 | 1 |
| HVRalt/4 | | | | | 8.71 | 1 |
| HVRalt/5 | | | | | 10.87 | 1 |
| HVRalt/6 | | | | | 11.33 | 1 |
| HVRalt/7 | | | | | 12.62 | 1 |
| HVRalt/8 | | | | | 13.17 | 1 |
| HVRalt/9 | | | | | 13.51 | 1 |
| HVRalt/10 | | | | | 13.85 | 1 |

| | | | | | | |
|---|---|---|---|---|---|---|
| **HVR, secondary structure from Huston, et al.[3] (29659– 29852)** | | | | | | |
| **HVRalt/1** | 194 | 1018079 | 25.06 | 23.99 ± 5.39 | 0 | 1 |
| HVRalt/2 | | | | | 0.91 | 1 |
| HVRalt/3 | | | | | 3.71 | 1 |
| HVRalt/4 | | | | | 3.78 | 1 |
| HVRalt/5 | | | | | 4.28 | 1 |
| HVRalt/6 | | | | | 6.65 | 1 |
| HVRalt/7 | | | | | 8.27 | 1 |
| HVRalt/8 | | | | | 8.53 | 1 |
| HVRalt/9 | | | | | 8.93 | 1 |
| HVRalt/10 | | | | | 9.72 | 1 |
| **3´ UTR BSL extended structure from Huston, et al.[3] (29543–29665; 29846–29876)** | | | | | | |
| **BSLext/1** | 158 | 1007602 | 25.81 | 24.60 ± 4.94 | 0 | 1 |
| BSLext/2 | | | | | 1.61 | 1 |
| BSLext/3 | | | | | 4.48 | 1 |
| BSLext/4 | | | | | 6.48 | 1 |
| BSLext/5 | | | | | 7.22 | 1 |
| BSLext/6 | | | | | 7.42 | 1 |
| BSLext/7 | | | | | 7.84 | 1 |
| BSLext/8 | | | | | 8.31 | 1 |
| BSLext/9 | | | | | 9.2 | 1 |
| BSLext/10 | | | | | 9.29 | 1 |
| **3´ UTR stem-loop II-like motif, secondary structure based on chemical reactivity from this study (29724–29773)** | | | | | | |
| **S2M/1** | 50 | 500000 | 2.56 | 5.76 ± 0.57 | 0 | 1 |
| S2M/2 | | | | | 0.29 | 2 |
| S2M/3 | | | | | 1.46 | 1 |
| S2M/4 | | | | | 1.57 | 1 |
| S2M/5 | | | | | 2.89 | 1 |
| S2M/6 | | | | | 3.04 | 2 |
| S2M/7 | | | | | 3.11 | 3 |
| S2M/8 | | | | | 3.88 | 1 |

| | | | | | | |
|---|---|---|---|---|---|---|
| S2M/9 | | | | | 4.02 | 2 |
| S2M/10 | | | | | 4.13 | 1 |
| **3´ UTR stem-loop II-like motif, secondary structure using chemical reactivity from Manfredonia, et al.[2] (29724–29773)** | | | | | | |
| **S2M/1** | 50 | 500000 | 2.68 | 5.86 ± 0.65 | 0 | 1 |
| S2M/2 | | | | | 1.14 | 3 |
| S2M/3 | | | | | 2.93 | 1 |
| S2M/4 | | | | | 4.6 | 1 |
| S2M/5 | | | | | 5.3 | 2 |
| S2M/6 | | | | | 5.32 | 1 |
| S2M/7 | | | | | 5.51 | 2 |
| S2M/8 | | | | | 5.8 | 1 |
| S2M/9 | | | | | 5.97 | 2 |
| S2M/10 | | | | | 6.38 | 1 |
| **3´ UTR stem-loop II-like motif, secondary structure using chemical reactivity from Huston, et al.[3] (29724–29773)** | | | | | | |
| **S2M/1** | 50 | 500000 | 2.74 | 5.91 ± 0.47 | 0 | 1 |
| S2M/2 | | | | | 0.19 | 1 |
| S2M/3 | | | | | 0.86 | 1 |
| S2M/4 | | | | | 0.94 | 1 |
| S2M/5 | | | | | 1.2 | 1 |
| S2M/6 | | | | | 1.27 | 1 |
| S2M/7 | | | | | 1.59 | 1 |
| S2M/8 | | | | | 1.61 | 1 |
| S2M/9 | | | | | 1.81 | 2 |
| S2M/10 | | | | | 2.23 | 1 |

[a]Mean pairwise all-heavy-atom RMSD between 10 lowest energy cluster centers.
[b]Predicted RMSD to true structure.
[c]Rosetta all-atom free energy gap of cluster's lowest energy model compared to lowest energy model discovered in run. REU = Rosetta energy units, calibrated so that 1.0 corresponds approximately to 1 $k_BT$.
[d]Number of models that appear in each cluster. Clustering was carried out on top 400 models ranked by Rosetta all-atom free energy, based on 5.0 Å threshold, except for small RNAs (SL1-4, SL6-7, s2m), where 2.0 Å threshold was applied.

**Table S4.** FARFAR2-Apo-Riboswitch models, extended data

| System/ cluster | Length | Models generated | Model convergence (Å)[a] | Predicted minimum RMSD (Å)[b] | E-gap to lowest energy model (REU)[c] | Cluster occupancy[d] | RMSD to experimental structure with ligand bound |
|---|---|---|---|---|---|---|---|
| **SAM-I riboswitch, RNA-Puzzle 4. PDB ID: 3V7E[11]** | | | | | | | |
| **SAM-I/1** | 126 | 5768 | 7.85 | 10.05 ± 2.55 | 0 | 99 | 2.52 |
| SAM-I/2 | | | | | 6.31 | 6 | 10.76 |
| SAM-I/3 | | | | | 8.03 | 211 | 8.23 |
| SAM-I/4 | | | | | 18.34 | 2806 | 9.68 |
| SAM-I/5 | | | | | 25.9 | 362 | 9.72 |
| SAM-I/6 | | | | | 26.8 | 22 | 14.83 |
| SAM-I/7 | | | | | 28.76 | 1 | 17.10 |
| SAM-I/8 | | | | | 29.06 | 282 | 11.90 |
| SAM-I/9 | | | | | 39.12 | 140 | 8.07 |
| SAM-I/10 | | | | | 39.77 | 588 | 11.33 |
| **SAM-I/IV riboswitch, RNA-Puzzle 8. PDB ID: 4L81[12]** | | | | | | | |
| **SAM-I/IV/1** | 96 | 33086 | 9.76 | 11.60 ± 2.88 | 0 | 4 | 8.83 |
| SAM-I/IV/2 | | | | | 1.77 | 7 | 11.18 |
| SAM-I/IV/3 | | | | | 2.75 | 19 | 10.98 |
| SAM-I/IV/4 | | | | | 3.65 | 15 | 5.23 |
| SAM-I/IV/5 | | | | | 5.16 | 4 | 9.62 |
| SAM-I/IV/6 | | | | | 5.99 | 2 | 12.25 |
| SAM-I/IV/7 | | | | | 6.84 | 1 | 7.69 |
| SAM-I/IV/8 | | | | | 8.78 | 2 | 7.44 |
| SAM-I/IV/9 | | | | | 9.07 | 15 | 5.36 |
| SAM-I/IV/10 | | | | | 9.21 | 1 | 14.13 |
| **SAM-IV riboswitch, RNA-Puzzle Unknown Rfam 15. PDB ID: 6UET[13]** | | | | | | | |
| **SAM-IV/1** | 119 | 10828 | 8.43 | 10.52 ± 2.39 | 0 | 4 | 7.61 |
| SAM-IV/2 | | | | | 2.16 | 6 | 3.70 |
| SAM-IV/3 | | | | | 8.44 | 3 | 6.04 |
| SAM-IV/4 | | | | | 10.13 | 8 | 11.37 |

| | | | | | | | |
|---|---|---|---|---|---|---|---|
| SAM-IV/5 | | | | | 10.15 | 1 | 9.55 |
| SAM-IV/6 | | | | | 12.88 | 3 | 7.03 |
| SAM-IV/7 | | | | | 14.4 | 3 | 8.68 |
| SAM-IV/8 | | | | | 14.72 | 1 | 8.51 |
| SAM-IV/9 | | | | | 17.51 | 2 | 8.01 |
| SAM-IV/10 | | | | | 17.94 | 2 | 6.28 |
| **Glycine riboswitch, RNA-Puzzle 3. PDB ID: 3OXE[14]** | | | | | | | |
| **Gly/1** | 84 | 33442 | 12.14 | 13.53 ± 2.13 | 0 | 1 | 17.96 |
| Gly/2 | | | | | 1.53 | 1 | 17.86 |
| Gly/3 | | | | | 3.87 | 1 | 21.51 |
| Gly/4 | | | | | 7.46 | 1 | 12.41 |
| Gly/5 | | | | | 7.76 | 1 | 14.54 |
| Gly/6 | | | | | 7.79 | 1 | 18.93 |
| Gly/7 | | | | | 8.2 | 1 | 16.66 |
| Gly/8 | | | | | 10.03 | 1 | 19.75 |
| Gly/9 | | | | | 10.71 | 1 | 15.60 |
| Gly/10 | | | | | 10.89 | 1 | 17.17 |
| **Cobalamin riboswitch, RNA-Puzzle 6. PDB ID: 4GXY[15]** | | | | | | | |
| **Cobalamin/1** | 158 | 28859 | 14.97 | 15.81 ± 2.42 | 0 | 1 | 20.39 |
| Cobalamin/2 | | | | | 0.47 | 1 | 13.08 |
| Cobalamin/3 | | | | | 2.41 | 1 | 24.45 |
| Cobalamin/4 | | | | | 4.14 | 1 | 19.51 |
| Cobalamin/5 | | | | | 4.84 | 1 | 17.94 |
| Cobalamin/6 | | | | | 6.1 | 1 | 24.51 |
| Cobalamin/7 | | | | | 7.48 | 1 | 19.98 |
| Cobalamin/8 | | | | | 9.07 | 1 | 25.93 |
| Cobalamin/9 | | | | | 9.74 | 1 | 19.14 |
| Cobalamin/10 | | | | | 11.56 | 1 | 16.42 |

| | | | | | | | |
|---|---|---|---|---|---|---|---|
| **5HT riboswitch, RNA-Puzzle 9. PDB ID: 5KPY[16]** | | | | | | | |
| **5HT/1** | 71 | 18660 | 7.89 | 10.08 ± 2.24 | 0 | 44 | 4.56 |
| 5HT/2 | | | | | 1.39 | 27 | 10.12 |
| 5HT/3 | | | | | 2.43 | 32 | 6.97 |
| 5HT/4 | | | | | 7.14 | 12 | 6.54 |
| 5HT/5 | | | | | 7.91 | 32 | 5.29 |
| 5HT/6 | | | | | 9.4 | 9 | 6.99 |
| 5HT/7 | | | | | 9.81 | 16 | 9.59 |
| 5HT/8 | | | | | 9.87 | 10 | 9.37 |
| 5HT/9 | | | | | 10 | 21 | 6.92 |
| 5HT/10 | | | | | 10.44 | 6 | 10.81 |
| **ydaO riboswitch, RNA-Puzzle 12. PDB ID: 4QLM[17]** | | | | | | | |
| **ydaO/1** | 117 | 35506 | 21.19 | 20.86 ± 2.43 | 0 | 1 | 23.35 |
| ydaO/2 | | | | | 1.66 | 1 | 16.62 |
| ydaO/3 | | | | | 4.02 | 1 | 17.26 |
| ydaO/4 | | | | | 5.53 | 1 | 18.62 |
| ydaO/5 | | | | | 7.2 | 1 | 14.28 |
| ydaO/6 | | | | | 7.21 | 1 | 18.13 |
| ydaO/7 | | | | | 7.4 | 1 | 16.90 |
| ydaO/8 | | | | | 7.75 | 1 | 13.32 |
| ydaO/9 | | | | | 8.13 | 1 | 16.23 |
| ydaO/10 | | | | | 9.54 | 1 | 19.93 |
| **ZMP riboswitch, RNA-Puzzle 13. PDB ID: 4XW7[18]** | | | | | | | |
| **ZMP/1** | 60 | 20297 | 10.45 | 12.16 ± 1.94 | 0 | 1 | 16.19 |
| ZMP/2 | | | | | 1.07 | 1 | 10.97 |
| ZMP/3 | | | | | 2.07 | 1 | 13.30 |
| ZMP/4 | | | | | 2.79 | 1 | 16.25 |
| ZMP/5 | | | | | 3.88 | 2 | 11.43 |
| ZMP/6 | | | | | 4.33 | 3 | 13.83 |
| ZMP/7 | | | | | 6.29 | 1 | 7.13 |
| ZMP/8 | | | | | 7.26 | 2 | 14.76 |
| ZMP/9 | | | | | 7.8 | 1 | 10.62 |
| ZMP/10 | | | | | 8.03 | 1 | 12.43 |
| **Glutamine riboswitch (bound), RNA-Puzzle 14. PDB ID: 5DDP[19]** | | | | | | | |

| | | | | | | | |
|---|---|---|---|---|---|---|---|
| **Gln (bound)/1** | 61 | 24531 | 11.97 | 13.39 ± 3.48 | 0 | 3 | 10.93 |
| Gln (bound)/2 | | | | | 1.34 | 2 | 11.71 |
| Gln (bound)/3 | | | | | 1.45 | 1 | 6.88 |
| Gln (bound)/4 | | | | | 5.31 | 5 | 11.96 |
| Gln (bound)/5 | | | | | 6.21 | 2 | 10.85 |
| Gln (bound)/6 | | | | | 7.27 | 1 | 10.28 |
| Gln (bound)/7 | | | | | 7.78 | 1 | 16.11 |
| Gln (bound)/8 | | | | | 8.28 | 8 | 11.33 |
| Gln (bound)/9 | | | | | 9.24 | 2 | 13.41 |
| Gln (bound)/10 | | | | | 9.64 | 1 | 10.02 |
| **Guanidinium riboswitch, RNA-Puzzle 21. PDB ID: 5NWQ[20]** | | | | | | | |
| **Guanidine/1** | 41 | 48146 | 9.26 | 11.19 ± 1.76 | 0 | 21 | Guanidine/1 |
| Guanidine/2 | | | | | 1.98 | 10 | Guanidine/2 |
| Guanidine/3 | | | | | 7.02 | 62 | Guanidine/3 |
| Guanidine/4 | | | | | 7.06 | 24 | Guanidine/4 |
| Guanidine/5 | | | | | 7.5 | 11 | Guanidine/5 |
| Guanidine/6 | | | | | 8.66 | 48 | Guanidine/6 |
| Guanidine/7 | | | | | 8.94 | 10 | Guanidine/7 |
| Guanidine/8 | | | | | 9.02 | 16 | Guanidine/8 |
| Guanidine/9 | | | | | 9.34 | 19 | Guanidine/9 |
| Guanidine/10 | | | | | 9.77 | 6 | Guanidine/10 |

[a]Mean pairwise all-heavy-atom RMSD between 10 lowest energy cluster centers.
[b]Predicted RMSD to true structure.
[c]Rosetta all-atom free energy gap of cluster's lowest energy model compared to lowest energy model discovered in run. REU = Rosetta energy units, calibrated so that 1.0 corresponds approximately to 1 $k_BT$.

[d]Number of models that appear in each cluster. Clustering was carried out on top 400 models ranked by Rosetta all-atom free energy, based on 5.0 Å threshold.

**Table S5**. Primers used in this study.

| Name | Purpose | Sequence (5′ to 3′) |
| --- | --- | --- |
| Extended 5′ UTR gBlock | DNA template for regions in the extended 5′ UTR | TTCTAATACGACTCACTATTATTAAAGGTTTATACCT TCCCAGGTAACAAACCAACCAACTTTCGATCTCTTGT AGATCTGTTCTCTAAACGAACTTTAAAATCTGTGTGG CTGTCACTCGGCTGCATGCTTAGTGCACTCACGCAGT ATAATTAATAACTAATTACTGTCGTTGACAGGACACG AGTAACTCGTCTATCTTCTGCAGGCTGCTTACGGTTT CGTCCGTGTTGCAGCCGATCATCAGCACATCTAGGTT TCGTCCGGGTGTGACCGAAAGGTAAGATGGAGAGCCT TGTCCCTGGTTTCAACGAGAAAACACACGTCCAACTC AGTTTGCCTGTTTTACAGGTTCGCGACGTGCTCGTAC GTGGCTTTGGAGACTCCGTGGAGGAGGTCTTATCAGA GGCACGTCAACATCTTAAAGATGGCACTTGTGGCTTA GTAGAAGTTGAAAAAGGCGTTTTGCCTCAACTTGAAC AGCCCTATGTGTTCATCAAACGTTCGGATGCTCGAACTG |
| 3′ UTR gBlock | DNA template for regions in the 3′ UTR | TTCTAATACGACTCACTATTTGAAACTCAAGCCTTAC CGCAGAGACAGAAGAAACAGCAAACTGTGACTCTTC TTCCTGCTGCAGATTTGGATGATTTCTCCAAACAATT GCAACAATCCATGAGCAGTGCTGACTCAACTCAGGC CTAAACTCATGCAGACCACACAAGGCAGATGGGCTA TATAAACGTTTTCGCTTTTCCGTTTACGATATATAGTC TACTCTTGTGCAGAATGAATTCTCGTAACTACATAGC ACAAGTAGATGTAGTTAACTTTAATCTCACATAGCAAT CTTTAATCAGTGTGTAACATTAGGGAGGACTTGAAAG AGCCACCACATTTTCACCGAGGCCACGCGGAGTACG ATCGAGTGTACAGTGAACAATGCTAGGGAGAGCTGC CTATATGGAAGAGCCCTAATGTGTAAAATTAATTTTA GTAGTGCTATCCCCATGTGATTTTAATAGCTTCTTAG GAGAATGAC |
| SL1-4 Forward Primer | PCR amplification for SL1-4 from 5′ UTR gBlock | TTCTAATACGACTCACTATTATTAAAGGTTTATACC |
| SL1-4 Reverse Primer | PCR amplification for SL1-4 from 5′ UTR gBlock | GTTGTTGTTGTTGTTTCTTTCAGTAATTAGTTATTAATT ATACTGCGTGAGTGC |
| Reverse complement of SL1-4 Forward Primer | PCR amplification for reverse complement of SL1-4 from 5′ UTR gBlock | TTCTAATACGACTCACTATTCAGTAATTAGTTATTAATT ATACTGCG |
| Reverse complement of SL1-4 Reverse Primer | PCR amplification for reverse complement of SL1-4 from 5′ UTR gBlock | ATTAAAGGTTTATACCTTCCCAGG |
| SL2-6_T7_CM-1F | DNA primer for PCR assembly of SL2-6 with 5’ and 3’ flanking “reference hairpins” | TTCTAATACGACTCACTATAGGGTCAGCGAGTAGCTG ACAACGATCTCTTGTAGATCTGTTCTCTAAACGAACTT TAAAATCTGTGTGGC |

| | | |
|---|---|---|
| SL2-6_T7_CM-2R | DNA primer for PCR assembly of SL2-6 with 5’ and 3’ flanking “reference hairpins” | GCAGCCGAGTGACAGCCACACAGATTT |
| SL2-6_T7_CM-3F | DNA primer for PCR assembly of SL2-6 with 5’ and 3’ flanking “reference hairpins” | ACTCGGCTGCATGCTTAGTGCACTCACGCAGTATAAT TAATAACTAATTACTGTCGTTGACAGGACACGAGTAAC TCGTCT |
| SL2-6_T7_CM-4R | DNA primer for PCR assembly of SL2-6 with 5’ and 3’ flanking “reference hairpins” | CCCGGACGAAACCTAGATGTGCTGATGATCGGCTGCA ACACGGACGAAACCGTAAGCAGCCTGCAGAAGATAGA CGAGTTACTCGTGTCCT |
| SL2-6_T7_CM-5F | DNA primer for PCR assembly of SL2-6 with 5’ and 3’ flanking “reference hairpins” | TTCGTCCGGGTGTGACCGAAAGGTAAGATGGAGAGCC TTGTCCCTGGTTTCAACGAGAAAACACACGTCCAACTC AGTTTGCC |
| SL2-6_T7_CM-6R | DNA primer for PCR assembly of SL2-6 with 5’ and 3’ flanking “reference hairpins” | GTTGTTGTTGTTGTTTCTTTGTGTCAGCTACTCGCTGAC TTCACGTCGCGAACCTGTAAAACAGGCAAACTGAGTTGG |
| Hyper-variable region Forward Primer | PCR amplification for hyper-variable region from 3′ UTR gBlock | TTCTAATACGACTCACTATTCTTTAATCTCACATAGCA ATCTTTAATC |
| Hyper-variable region Reverse Primer | PCR amplification for hyper-variable region from 3′ UTR gBlock | GTTGTTGTTGTTGTTTCTTTTATTAAAATCACATGGG GATAGCACTAC |
| FAM-A20-Tail2 | RNA extraction and cDNA labeling | /56-FAM/AAAAAAAAAAAAAAAAAAAAGTTGTTGTTGTTGTTTCTTT |
| RTB000 | RT primer for no-modification sample | /56-FAM/AATGATACGGCGACCACCGAGATCTACACTC TTTCCCTACACGACGCTCTTCCGATCTACCAGGCGCT GGGTTGTTGTTGTTGTTTCTTT |
| RTB001 | RT primer for 1M7 modification sample | /56-FAM/AATGATACGGCGACCACCGAGATCTACACTCTTT CCCTACACGACGCTCTTCCGATCTGAGGCCTTGG CCGTTGTTGTTGTTGTTTCTTT |
| pA-Adapt-Bp | Linker ligated for Illumina sequencing | /5Phos/AGATCGGAAGAGCGGTTCAGCAGGAATGCC GAGACCGATCTCGTATGCCGTCTTCTGCTTG/3Phos/ |
| Eterna Construct 1, ID 9850089 | Eterna construct for chemical mapping of genomic positions 45-107 | TTCTAATACGACTCACTATTGGAAAGATCTCTTGTAG ATCTGTTCTCTAAACGAACTTTAAAATCTGTGTGGCT GTCACTCGGCTGCAGCAGACGTTCGCGTCTGCAAAA GAAACAACAACAACAAC |
| Eterna Construct 2, ID 9849692 | Eterna construct for chemical mapping of genomic positions 76-136 | TTCTAATACGACTCACTATTGGAAATTTTAAAATCTGT GTGGCTGTCACTCGGCTGCATGCTTAGTGCACTCACGCAG TATAATTAAACGTGGGCTTCGGCTCGCGAAAAGAAAC AACAACAACAAC |
| Eterna Construct 3, ID 9872605 | Eterna construct for chemical mapping of genomic positions 172-234 | TTCTAATACGACTCACTATTGGAAATCGTCTATCTT CTGCAGGCTGCTTACGGTTTCGTCCGTGTTGCAGC CGATCATCAGCACATCTCCGAAGCTTCGGCTTCG GAAAAGAAACAACAACAACAAC |

| | | |
|---|---|---|
| Eterna Construct 4, ID 9849874 | Eterna construct for chemical mapping of genomic positions 172-233 | TTCTAATACGACTCACTATTGGAAATCGTCTAT CTTCTGCAGGCTGCTTACGGTTTCGTCCGTG TTGCAGCCGATCATCAGCACATCAGCAGCTGT TCGCAGCTGCAAAAGAAACAACAACAACAAC |
| Eterna Construct 5, ID 9872974 | Eterna construct for chemical mapping of genomic positions 217-277 | TTCTAATACGACTCACTATTGGAAACCGATCAT CAGCACATCTAGGTTTCGTCCGGGTGTGACCG AAAGGTAAGATGGAGAGCATTAAGCGGGTCTT CGGGCCTGCAAAAGAAACAACAACAACAAC |
| Eterna Construct 6, ID 9872770 | Eterna construct for chemical mapping of genomic positions 13472-13529 | TTCTAATACGACTCACTATTGGAAAGATTTGCGG TGTAAGTGCAGCCCGTCTTACACCGTGCGGCAC AGGCACTAGTACTGATGTAAACGGCGGCTTCGG TCGTTGAAAAGAAACAACAACAACAAC |
| Eterna Construct 7, ID 9873087 | Eterna construct for chemical mapping of genomic positions 29720-29776 | TTCTAATACGACTCACTATTGGAAAAGAGACACC ACATTTTCACCGAGGCCACGCGGAGTACGATCG AGTGTACAGTGAACAATGCTACGATACGTTCGC GTATCGAAAAGAAACAACAACAACAAC |
